# Human land-use change drives co-occurrence of ecologically similar avian aerial insectivores in Southeast Asia

**DOI:** 10.64898/2026.05.20.726292

**Authors:** Alexis M. Garvin, Samantha S. Sudoko, Nurhartini K. Yahya, Nur Alwanie Maruji, Rayzigerson R. Chai, Karim A. bin Dakog, Benoit Goossens, Jamie M. Kass, Elizabeth S.C. Scordato

**Affiliations:** Department of Biological Sciences, California State Polytechnic University, Pomona, CA, USA, 91768; Department of Evolution, Ecology, and Organismal Biology, University of California, Riverside, CA, USA, 92521; Danau Girang Field Centre, c/o Sabah Wildlife Department, Wisma MUIS, Block B 5^th^ Floor, 88100 Kota Kinabalu, Sabah, Malaysia; Wildlife Health, Genetic and Forensic Laboratory, c/o Sabah Wildlife Department, Wisma MUIS, Block B 5^th^ Floor, 88100 Kota Kinabalu, Sabah, Malaysia; Sabah Wildlife Department, Wisma MUIS, Block B 5^th^ Floor, 88100 Kota Kinabalu, Sabah, Malaysia; Organisms and Environment Division, School of Biosciences, Cardiff University, Sir Martin Evans Building, Museum Avenue, Cardiff CF10 3AX, UK; Graduate School of Life Sciences, Tohoku University, Sendai, Miyagi, Japan

**Author notes:** Address correspondence to Elizabeth Scordato.

**Keywords:** Joint species distribution model, human land-use change, niche partitioning, biotic interactions, aerial insectivores

## Abstract

**Aim:** Human land-use change contributes to biodiversity declines, but also creates new niches that facilitate novel biotic interactions. These interactions can reshape ecological communities and ecosystem function, yet remain poorly understood. Swiftlets and swallows in Southeast Asia present a classic example: coexistence is facilitated by fine-scale diet partitioning, with population sizes historically limited by available nesting substrates. However, several species now nest on manmade structures, particularly “nest farms” built to harvest edible swiftlet nests. We evaluated whether land-use change, especially the spread of nest farms, is leading to breakdowns in niche partitioning and increased competition among six sympatric swiftlets and swallows.

**Location:** Northern Borneo

**Methods:** We calculated geographic niche overlap using species distribution models (SDMs) with different environmental predictors, hypothesizing greater overlap when land-use variables were included. We then implemented joint species distribution models (JSDMs) to partition shared environmental responses from potential biotic interactions, predicting that competition would emerge as negative residual correlations. We used sightings from citizen-science datasets and structured surveys to evaluate the influence of climate, land-use, nest farms, morphology, and foraging behavior on species occurrences.

**Results:** SDMs that included land-use variables showed high niche overlap, suggesting that human activity homogenizes niches. The optimal JSDM, based on structured survey data, identified distance to nest farms as the strongest predictor of occurrence for all species, with species showing both positive and negative responses. Morphology and behavior had small effects, and residual correlations were weak, indicating limited unexplained biotic interactions.

**Main conclusions:** Human activity, through the creation of artificial nesting sites, broadly drives co-occurrence of swallows and swiftlets across our study region. These effects appear to operate primarily through environmental filtering rather than direct competition. Our findings reveal substantial and complex impacts of land-use change and anthropogenic nest sites on the distribution and composition of aerial insectivore communities.

## Introduction

Human land-use change can dramatically alter resource distributions within the boundaries of a species’ climatic niche (Boivin et al., 2016; Ellis et al.; 2021). Reductions in suitable habitat may force species into closer proximity and affect biotic interactions, for example by increasing competition (Rowe 2009; Lee et al., 2024). Human land-use can also create new niches within a species’ existing range or facilitate movement into previously inaccessible areas, thereby introducing novel interspecific interactions (Essl et al., 2019; Diggins et al., 2023) or increasing the frequency of once-rare interactions (Gilbert et al., 2022; Sáenz-Jiménez et al., 2020). This in turn may disrupt longstanding dietary, temporal, spatial, or behavioral niche partitioning (Acevedo & Casinello 2009; Wang et al. 2020), leading to altered community dynamics (Sévêque et al. 2020; Parsons et al. 2019). However, quantifying species interactions in wild populations remains challenging (Fayle et al. 2015; Siepielski & McPeek 2010), and the effects of human land-use change on biotic interactions remain poorly understood.

To address these challenges, researchers have developed methods to infer interspecific interactions from patterns of co-occurrence. One such approach is modeling geographic or ecological niche overlap, which can identify instances of potential competitive exclusion (e.g. Novella-Fernandez et al. 2021, Nesbit et al. 2023, Oeser et al. 2025). Comparing niche overlap among models built with different predictor sets (e.g., climate and land-use, Ramirez et al. 2025), can further generate hypotheses about how land-use change may alter biotic interactions; for example, niche overlap among species exploiting anthropogenic environments may reveal emergent competitive interactions. However, assessing potential niche overlap alone does not account for shared environmental responses, shared evolutionary history, or functional variation between species (Murray et al. 2023). Failure to account for these factors limits inference, as observed niche overlap may reflect environmental filtering or shared responses to unmeasured habitat features rather than true biotic interactions.

Joint species distribution models (JSDMs) partially overcome these limitations by simultaneously modeling multiple species’ responses to shared environmental predictors. JSDMs can incorporate phylogenetic and functional trait data, enabling assessment of whether species with similar traits or evolutionary histories are more or less likely to co-occur (Ovaskainen et al. 2017). They also use latent variables to model co-occurrence variance that remains after controlling for environmental covariates. These residual correlations can reflect potential biotic interactions, such as competition (negative correlations) or facilitation (positive correlations, Warton, 2015). However, it remains challenging to disentangle biotic interactions from unmeasured environmental variables, both of which can generate residual associations in JSDMs (Poggiato 2021). This is a particular concern when occurrence is influenced by landscape features that are species-specific, difficult to measure, or are not reflected in widely available databases. Therefore, while JSDMs offer a framework for exploring potential biotic interactions, careful interpretation is essential for accurate ecological inference (Blanchet et al., 2020).

An additional challenge to JSDMs is the type of occurrence data used for modeling. JSDMs require absence data, unlike single-species SDMs that can use pseudoabsence or background data, and reliable absence data can be limited. Structured surveys, while providing high-quality data, are often narrow in spatial and temporal coverage. To address these gaps, semi-structured citizen-science data have become a common component of SDM studies (Feldman et al. 2021; Johnston et al. 2018). However, data collection is opportunistic, which introduces biases such as uneven sampling effort and spatial sampling bias (Gorleri et al. 2022; Isaac et al. 2015). While some studies have found that combining citizen-science data with structured surveys can improve the predictive accuracy of SDMs (Robinson et al., 2020), there are few examples using citizen-science data in JSDMs.

In this study, we combined SDM-based measurements of geographic niche overlap and JSDMs to assess how human land-use change may influence co-occurrence of aerial insectivorous birds in Southeast Asia (Figure 1). This region has transformed over the past several decades, with forests and rural areas converted to oil palm plantations and urban developments (Koh & Wilcove, 2007). Aerial insectivores are particularly vulnerable to these changes, as intensively managed agriculture can reduce arthropod food (Ghazali et al. 2021). We focused on four species of swiftlets (genera *Aerodramus* and *Collocalia*) and two species of swallows (genus *Hirundo*), all of which are widely distributed, ecologically similar, and co-occur in sympatry across much of their ranges. Coexistence is thought to stem primarily from fine-scale dietary niche partitioning (Waugh & Hails 1983; Lourie & Tompkins 2008; Chan & Goh 2019). They also partition nest sites, with swallows and some swiftlets nesting on cliff faces and near the mouths of caves, while other swiftlets nest in cave interiors (Cranbrook et al. 2013). Population sizes were likely historically limited by nest site availability (Ramirez et al. 2025). However, some species (*H. javanica, C. affinis*) now nest extensively on artificial structures like bridges and buildings. Furthermore, the nests of Black-nest swiftlets (*A. maximus*) and White-nest swiftlets (*A. fuciphagus*) are culinary delicacies in parts of East Asia (Lee et al. 2021), which has led to the collapse of many colonies due to long-term overharvesting. Recently, however, demand for nests has driven the construction of tens of thousands of “nest farms” throughout the species’ ranges, potentially fueling population size increases and expansions into previously unoccupied regions (Cranbrook et al. 2013). Such large-scale modification of nest-site availability may disrupt historic niche partitioning and generate novel biotic interactions.

**Figure 1.**
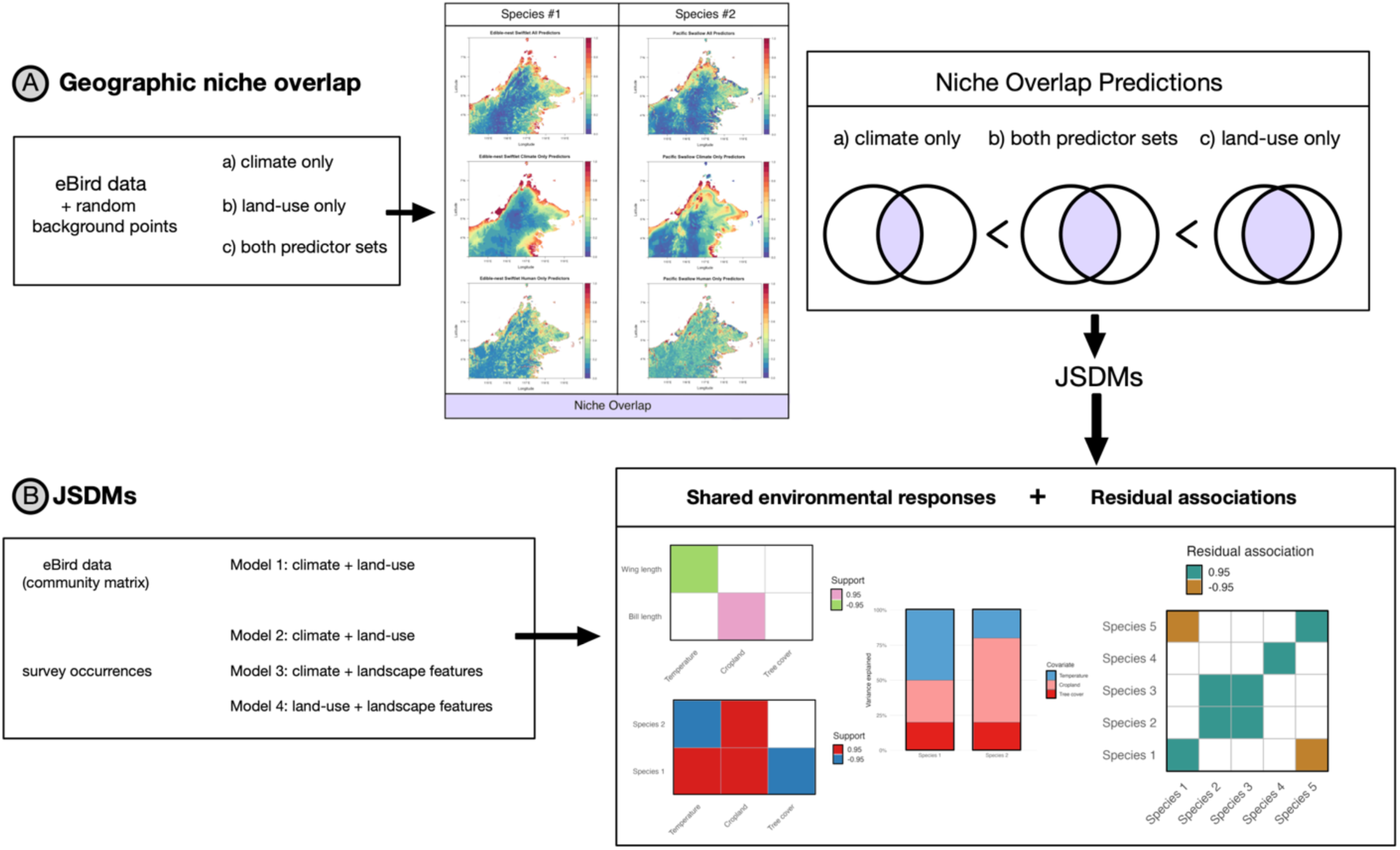
Overview of study design and analytical framework. We used SDM-based measures of niche overlap and JSDMs to assess how human land-use change influences patterns of co-occurrence among aerial insectivorous birds in Borneo. (A) We first compared potential geographic niche overlap between focal species under three environmental predictor sets: (a) climate data only, (b) land use data only, and (c) both predictor sets, hypothesizing that models incorporating land-use data would predict greater niche overlap. We then used JSDMs (B) to test whether models fit with citizen-science data produced comparable results to those built from structured surveys, and whether species co-occurrence patterns were best explained by climate, land use, or landscape features (nest farms and distance to water). We partitioned shared environmental responses from residual associations that could reflect biotic interactions.

We hypothesized that human land-use change, and particularly the use of anthropogenic structures for nesting, has homogenized historic niches and promoted increased co-occurrence among ecologically similar species, potentially intensifying competition. To evaluate this, we first quantified potential geographic niche overlap among our focal species using citizen-science-based occurrence records and three sets of predictor variables: (a) climate variables only, (b) human land-use variables only, and (c) all variables. If human land-use homogenizes niches, we expected models incorporating land-use variables to predict greater overlap among species compared to climate-only models (Fig. 1A). Next, we used a series of JSDMs to examine predictors of co-occurrence and disentangle potential biotic interactions from shared environmental responses. We assessed whether models fit with citizen-science data or structured survey data generated better model fits; whether the inclusion of landscape features relevant to our focal species, like nest farms, improved explanatory power over more general variables derived from remote sensing; and whether morphological or behavioral traits explained additional variation (Fig. 1B–C). If human land-use change is altering competition, we expected to find negative residual associations after accounting for shared environmental responses, morphology, and behavior.

## Methods

### Study area

Our study was conducted in Sabah, a Malaysian state on the island of Borneo (Figure 2A). Sabah’s lowland forests have been widely replaced by oil palm (Koh & Wilcove 2007; Gaveau et al. 2022), and urban development is increasing (Jakobsen et al., 2007). Sabah is ideal for our objectives because in addition to experiencing rapid land-use transformation it also supports a high density of sympatric swiftlets and swallows. The Pacific swallow (*H. javanica*), White-nest swiftlet (*A. fuciphagus*), Black-nest swiftlet (*A. maxima*), Mossy-nest swiftlet (*A. salangana*), and Plume-toed swiftlet (*C. esculenta*) are all sympatric residents co-occurring across broad geographic areas, while Barn swallows (*H. rustica*) are seasonal migrants that overwinter from October to May.

**Figure 2.**
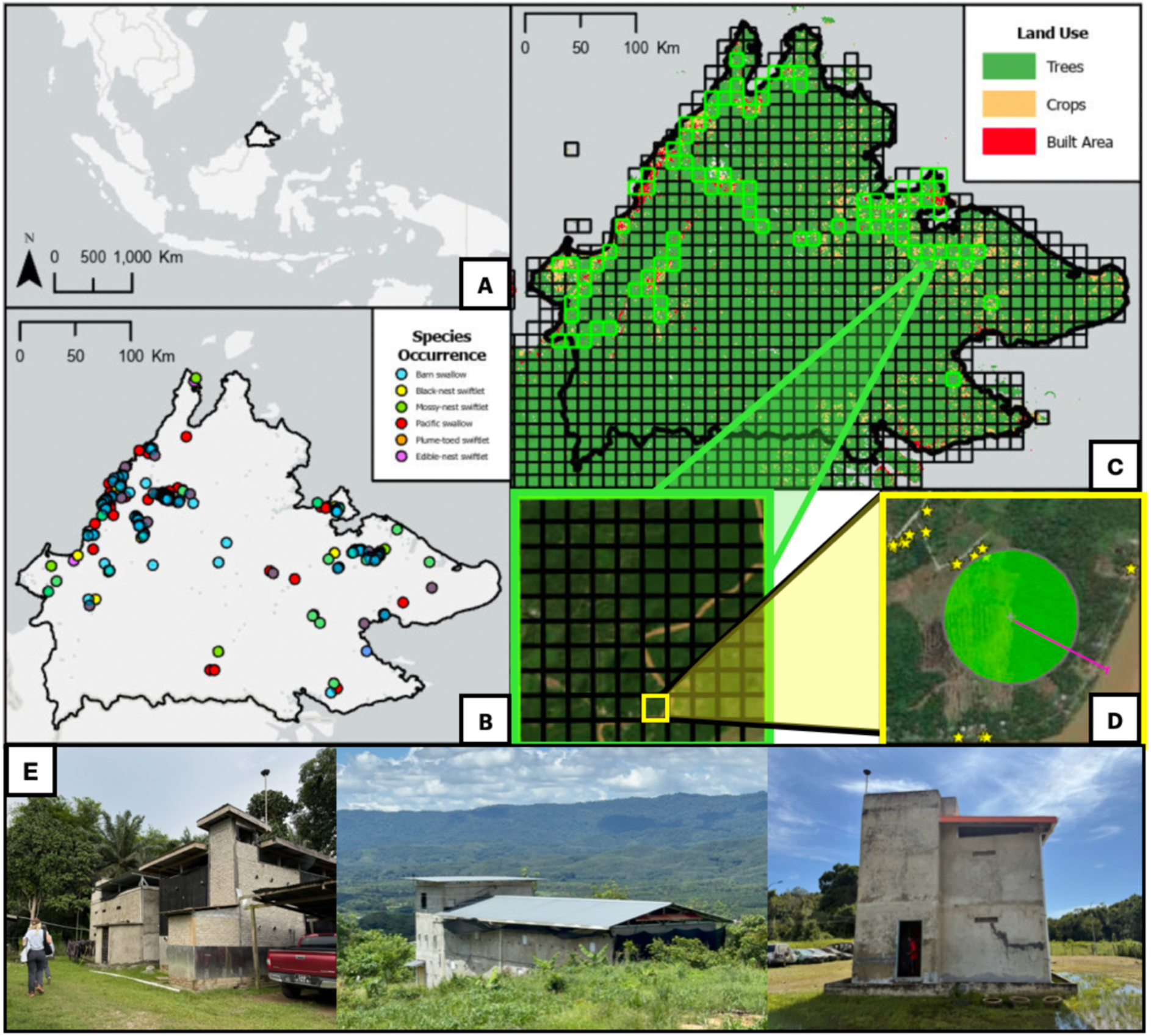
Study area and survey design. (A) Study region in Southeast Asia, focusing on the state of Sabah (outlined in black) in northern Borneo. (B) Species occurrence records from eBird for each focal species; some species overlap in the same location. (C) Distribution of 10 km² grid cells across the study extent. Neon green indicates cells containing 1 km² surveyed grid cells. (D) 1 km² survey grids, where green buffers show surveyed areas, yellow stars mark recorded nest farm locations, and the pink line illustrates an example post-processing metric used to calculate nearest distance to water. (E) Examples of nest farms, illustrating their distinctive architecture.

### Citizen-science occurrence data

We assembled two occurrence datasets: citizen-science records and structured survey data. For the former, we downloaded records from the eBird database (Sullivan et al. 2009) from 2015 to 2024 for each of our six focal species. To control for sampling bias, we used the R package *auk* (Strimas-Mackey et al. 2018) to filter occurrence data according to best practices (Johnston et al. 2021, Supplemental Material). Only complete checklists (i.e., those that reported all species detected during surveys) were retained. SDMs used only occurrence records, with pseudoabsences generated in the models (Supplemental Material). For JSDMs, we generated presence and absence data for each species. Presence locations were defined as 1km^2^ grid cells with at least one occurrence record of the focal species across all available checklists. Absence locations were grid cells where complete eBird checklists were recorded but the focal species was not observed. This yielded a community presence/absence matrix for our six species with 1,430 unique locations (Figure 2B, Supplemental Material).

### Structured occurrence surveys

We conducted systematic surveys of swallows and swiftlets across Sabah using a stratified sampling design to balance survey effort across land cover types. We overlaid a 1 km^2^ grid on our study area (bounding box: 114–120°E, 4–7°N) in ArcGIS Pro (Esri, 2024) and classified each grid cell (n= 124,300) by its dominant land cover type (urban area, cropland, or forest, Figure 2C). Within each grid cell, we circumscribed a 200m radius buffer around the centroid for surveying (Figure 2D). Much of the land in Sabah is forested and inaccessible, so we surveyed cells that were accessible by road while attempting balanced coverage across land-cover categories (urban area: n = 46, cropland: n = 67, forest: n = 40). To minimize spatial autocorrelation and ensure independence between sampling locations, we surveyed a maximum of four 1 km^2^ grid cells within any 10 km^2^ cell, and spaced surveyed cells at least 2 km apart. From 2024-2025, grid cells were surveyed once between May and June to restrict sampling to the dry season when many birds were incubating eggs or had chicks. This improved detectability and logistical feasibility. Swiftlets and Pacific swallows are year-round residents, and local densities are not expected to vary substantially throughout the year. Daily movements typically range from 1 to 16 km (Burhanuddin et al., 2017), and birds roost on their nests each night.

Surveys were conducted for 15 minutes within each 200m buffer between sunrise and 1100h and from 1600h to sunset, periods when birds forage lower in the air column and closer to vegetation. Three observers (A.M.G., S.S.S,, E.S.C.S.) systematically walked across the buffer to maximize coverage, recording all observations of the focal species. For grid cells in which large portions of the buffer were inaccessible (e.g., due to forest cover), observers walked the length of the roads and scanned the tops of trees with binoculars. Individuals flying above 100m were not recorded. As swiftlets and swallows forage above vegetation and road surfaces and rarely under the forest canopy, there was high confidence that most present birds were able to be identified. Black-nest and White-nest swiftlets are morphologically very similar. We therefore required all three observers to agree on identifications. A large proportion of our sightings occurred close to nest farms, and where possible we asked farm owners to confirm which species were there; these were typically consistent with our identifications. Following each survey, we searched for nests and documented the presence and density of nest farms within or adjacent to the survey buffer. We also recorded all nest farms visible while traveling between survey sites and through grid cells. Nest farms typically have distinctive and conspicuous architecture (Figure 2E), and while it was not possible to exhaustively sample every 10km^2^ grid cell, we have confidence that our data capture relative differences in nest farm abundance.

### Environmental predictors

To identify environmental predictors associated with occurrences, we downloaded 17 bioclimatic variables from CHELSA v2.1 (Karger et al. 2023) and 11 land-use variables from FAO-GLC (FAO, 2015) and coarsened them to 10 km^2^ resolution using bilinear interpolation for continuous variables and nearest neighbor distances for categorical variables. While conducting field sampling, we noticed that the GLC often misclassified cropland (typically oil palm) as tree cover. We therefore manually reclassified areas of clearly identifiable cropland by referencing high-resolution World Imagery basemaps (World Imagery, Esri). Additionally, we calculated the following landscape features: distance to the nearest nest farm, the number of nest farms within a 10-km radius (derived from our field surveys, Figure 2D-E), and distance to the nearest water body (coastline or river), calculated in ArcGIS (Figure 2D).

To reduce model overfitting, we selected a subset of uncorrelated variables from our original set (Merow et al. 2013). We generated a Pearson correlation matrix and chose one variable from each pair with |*r*| > 0.7, prioritizing more general measurements. The final variable set included mean annual temperature, mean annual precipitation, and isothermality as climate variables; artificial surfaces, cropland, and non-crop tree cover as land-use variables; and distance to water, distance to nearest nest farm, and number of nest farms within 10 km as landscape features (Supplemental Material).

### Species distribution models and geographic niche overlap

To assess the extent of potential geographic niche overlap among species, we built SDMs using the eBird occurrence records (Supplemental Material). Briefly, we built three sets of SDMs for each species reflecting different land use scenarios: models that included only climate variables (“climate-only”), models with only land-use variables (“land-use only”), and models with all variables (“both sets”) following Ramirez et al. (2025). Models were built using the Maxent algorithm (v3.4.4, Phillips et al., 2017) and tuned for optimal complexity in the R package ‘ENMeval’ (v2.0.4; Kass et al., 2021). We employed the “block” method for spatial cross-validation (Muscarella et al., 2014) and chose the model with the best combination of tuning parameters using the Continuous Boyce Index (CBI, Hirzel et al., 2006). We compared models built with different predictor sets using AICc and null models. For all species pairs and each variable set, we estimated geographic overlap of SDM predictions with the Schoener’s D statistic, which calculates similarity between two continuous rasters, where 0 is completely different and 1 is identical (Warren et al. 2008). We then assessed whether extent of geographic niche overlap differed among variable sets using a one-way ANOVA followed by Tukey HSD post-hoc tests.

### Morphological and behavioral trait data

We assembled morphological and behavioral data for each focal species. Morphological data were obtained from AVONET (Tobias et al. 2022). We chose functional traits expected to reflect habitat use (Violle 2007): bill depth and culmen length as indicators of foraging niche, wing and tail length for dispersal ability and flight performance, and body mass for biomass. We collected data on foraging traits from our field surveys, specifically group size and minimum, maximum, and modal foraging heights (estimated with a rangefinder; Nikon Forestry Pro 2), as these behaviors may reflect niche partitioning, particularly in mixed-species flocks (Schoener 1971; Naikatini et al. 2022).

### Phylogenetic relationships

To control for potential niche similarity among closely related species, we incorporated a phylogenetic random effect in our JSDMs. We downloaded 1000 trees from Vertlife phylogeny subsets (https://vertlife.org/phylosubsets/; Jetz et al. 2012) for all sequenced *Aerodramus, Collocalia,* and *Hirundo* species, constructed a consensus maximum-likelihood tree using the ‘ape’ package in R (v5.0; Paradis & Schliep, 2019), and pruned the tree to include only the focal species in our study (Supplemental Material).

### JSDM analysis

We used the Hierarchical Modeling of Species Communities (HMSC) framework (Ovaskainen 2017; Ovaskainen & Abrego 2020) to run probit regression models for the six species. We constructed four sets of JSDMs using different sources of occurrence data and combinations of predictor variables (Figure 1, Supplemental Material). To explore differences in model explanatory power for different occurrence data types, Models 1 and 2 both used climate and land-use predictor variables but different occurrence data: eBird occurrence records (same as for the SDMs) for Model 1, and structured survey data for Model 2. To explore differences in model explanatory power for different predictor variable types, Models 3 and 4 both used structured survey data but different predictor variables: climate variables and landscape features for Model 3, and land-use variables and landscape features for Model 4. All models had the same number of predictors (n = 6). For each model, we first built a JSDM that included only environmental predictors and spatial random effects, specified as the identity of each surveyed 1 km^2^ grid cell nested within its corresponding 10 km^2^ grid cell to account for spatial autocorrelation (Tikhonov 2020; Ovaskainen et al. 2017). We then compared this model to one that incorporated a phylogenetic tree and trait data (either morphometric or behavioral) and assessed whether this improved model fit (Figure 1, Supplemental Material). For Models 2, 3, and 4 we excluded the Barn swallow, which was not present during our surveys, as well as the Mossy-nest swiftlet, for which we only had three confirmed sightings among 155 surveys (Supplemental Material).

### JSDM fit and performance

JSDMs were fit using the R package ‘hmsc**’** (Tikhonov 2020), with four Markov Chain Monte Carlo (MCMC) chains, each run for 250,000 iterations. We discarded the first 125,000 iterations as burn-in and used a thinning interval of 1000, yielding 125 posterior samples per chain and 500 samples for posterior inference. We assessed MCMC convergence using the potential scale reduction factor (PSRF). We considered PSRF less than 1.2 to indicate good convergence (Gelman & Rubin 1992). Separate PSRF values were computed for β parameters (environmental covariates), Γ parameters (traits), ρ parameter (phylogenetic signal) and Ω parameters (residuals) for each model.

We assessed the in-sample explanatory power of each model using three complementary statistics: Area Under the Receiving Operating Characteristic Curve (AUC), which reflects the model’s ability to discriminate between presence and absence data; root mean square error (RMSE), which measures the model’s predictive accuracy by comparing the predicted values to observed occurrences; and Tjur’s R^2^, defined as the difference between the mean predicted occurrence probability for the occupied grid cells and the mean predicted occurrence probability for the unoccupied grid cells.

To compare the relative predictive power of different models, we used random four-fold cross-validation for Tjur’s R^2^ and AUC. We also calculated the Watanabe-Akaike Information Criterion (WAIC), which balances model fit with model complexity. To choose the best models, we first assessed ΔWAIC. When ΔWAIC < 2, we chose the model with the highest cross-validated Tjur’s R^2^. We additionally inspected cross-validated AUC to verify that the selected model also performed well at distinguishing presences from absences.

### Variance partitioning and parameter estimates

We used variance partitioning to quantify the proportion of variance in species occurrences (based on Tjur’s R2) explained by each environmental predictor and the spatial random effect (Ovaskainen & Abrego, 2020). To assess the direction and magnitude of these associations, we extracted parameter estimates for environmental predictors (β), traits (Γ), and residual associations (Ω). To visualize how community composition and species occurrence probabilities varied along environmental gradients, we constructed partial dependence plots based on posterior predictions for variables with high explanatory power in the optimal model.

## Results

### Geographic niche overlap

The SDMs with both climate and human land-use predictors had the lowest AICc for all species except the Barn swallow, with selected feature classes and regularization multipliers indicating relatively simple to moderately complex models (Supplemental Material). Niche overlap values were uniformly high (mean= 0.80 ± 0.05) yet differed significantly among predictor variable sets (*F*_(2, 42)_ = 8.44, *p* < .001), with the highest overlap values for models with only human land-use predictors (Tables S5-S7, Supplemental Material). The highest overlap was Barn Swallow – Pacific Swallow and Pacific Swallow – Plume-toed Swiftlet, which are the three species most closely associated with human-modified environments. These results support the hypothesis that land-use change may homogenize available habitats and increase geographic niche overlap. However, the relatively lower overlap in the climate-only and combined models suggests that some environmental filtering based on climate may persist among species.

### JSDMS Model Fit and Performance

All JSDMs demonstrated satisfactory convergence (Supplemental Material). WAIC varied by <2 among models, so we used the average Tjur’s R² and AUC from cross-validation (CV) to select an optimal model. We refer to parameters with Bayesian posterior support of ≥ 0.95 as strongly supported and those with posterior support between 0.80–0.95 as moderately supported. Hmsc computes in-sample metrics separately for each species. We report the mean value across species for each model in Table 1 and the species-specific values in Supplemental Material.

**Table 1.**
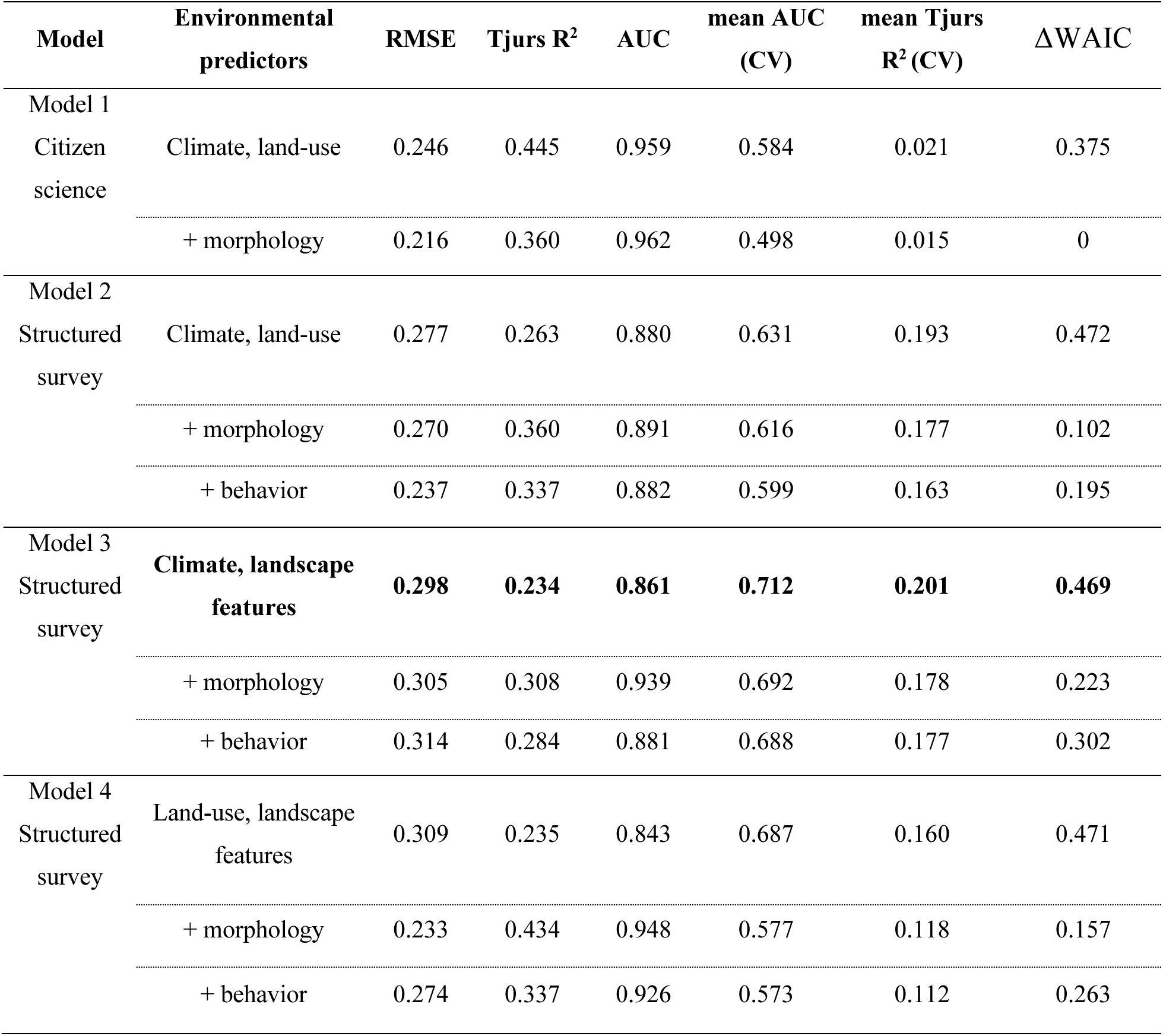
Comparison of model performance across four JSDMs. Models differed by source of occurrence records and by environmental predictors (n = 6 predictors for all models). Model 1 used citizen-science occurrences with climate and land-use predictors. Model 2 used structured survey occurrences with climate and land-use predictors. Models 3 and 4 used structured survey occurrences with landscape features and either climate (Model 3) or land-use predictors (Model 4). Columns indicate in-sample and out-of-sample (cross-validation: CV) performance metrics. Lower RMSE values indicate better in-sample fit. Higher Tjur’s R² values indicate better discrimination between presences and absences. AUC reflects classification accuracy, with values closer to 1.0 indicating better performance. Four-fold cross-validated metrics (mean CV AUC, mean CV Tjur’s R²) assess predictive performance and generalizability. For WAIC, lower values indicate better model performance after penalizing model complexity. Model 1 includes two species not present in surveys (Barn swallows and Mossy-nest swiftlets) and therefore does not include behavioral data. The best overall model is boldfaced.

### Model 1: Citizen-science-based occurrence records with climate and land-use predictors

Model 1 had the highest mean AUC and a relatively low RMSE when evaluating on in-sample data (AUC = 0.959; RMSE = 0.246), but cross-validation revealed poor predictive performance for withheld data (mean AUC = 0.584; mean Tjur’s R² = 0.021, Table 1). This reflects severe overfitting, likely due to spatial sampling bias in the unstructured survey data. Consistent with low predictive power, variance partitioning showed that the spatial random effect drove most of the variation (Figure 3A, Table 2). Several environmental covariates had strong posterior support (Figure 4A), but parameter estimates (β) were very small (Table 2, Supplemental Material). The residual correlation matrix revealed nearly perfect pairwise correlations (Supplemental Material), indicating that after accounting for environmental variation the remaining unexplained co-occurrence among species was extremely high, likely due to unmeasured covariates or spatial sampling bias.

**Figure 3.**
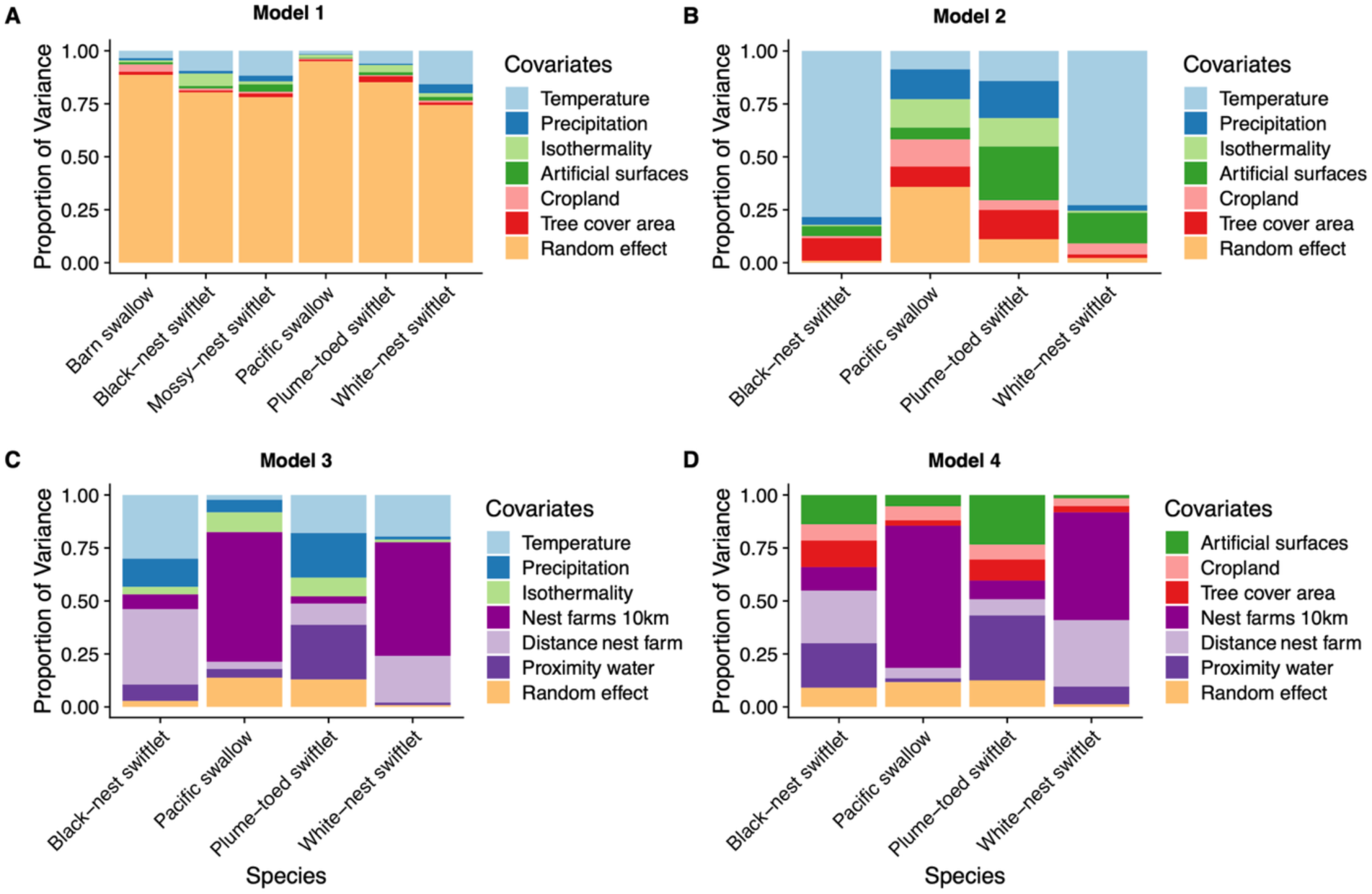
Variance partitioning across four JSDMs. Each bar represents the proportion of variance in species occurrences explained by each environmental covariate and the spatial random effects in the model. Covariates included climate variables, land-use predictors, and landscape features depending on the model. A) Model 1 used citizen-science occurrence records with climate and land-use predictors. B) Model 2 used structured survey records with climate and land-use predictors. Models 3 and 4 used structured survey data with landscape features and either climate (C) or land-use (D). Model 3 was the best fit to the data. Model 1 (A) was constructed using citizen-science occurrence records and includes the Barn swallow and Mossy-nest swiftlet. These species were absent from structured surveys (Models 2-4, B-D).

**Figure 4.**
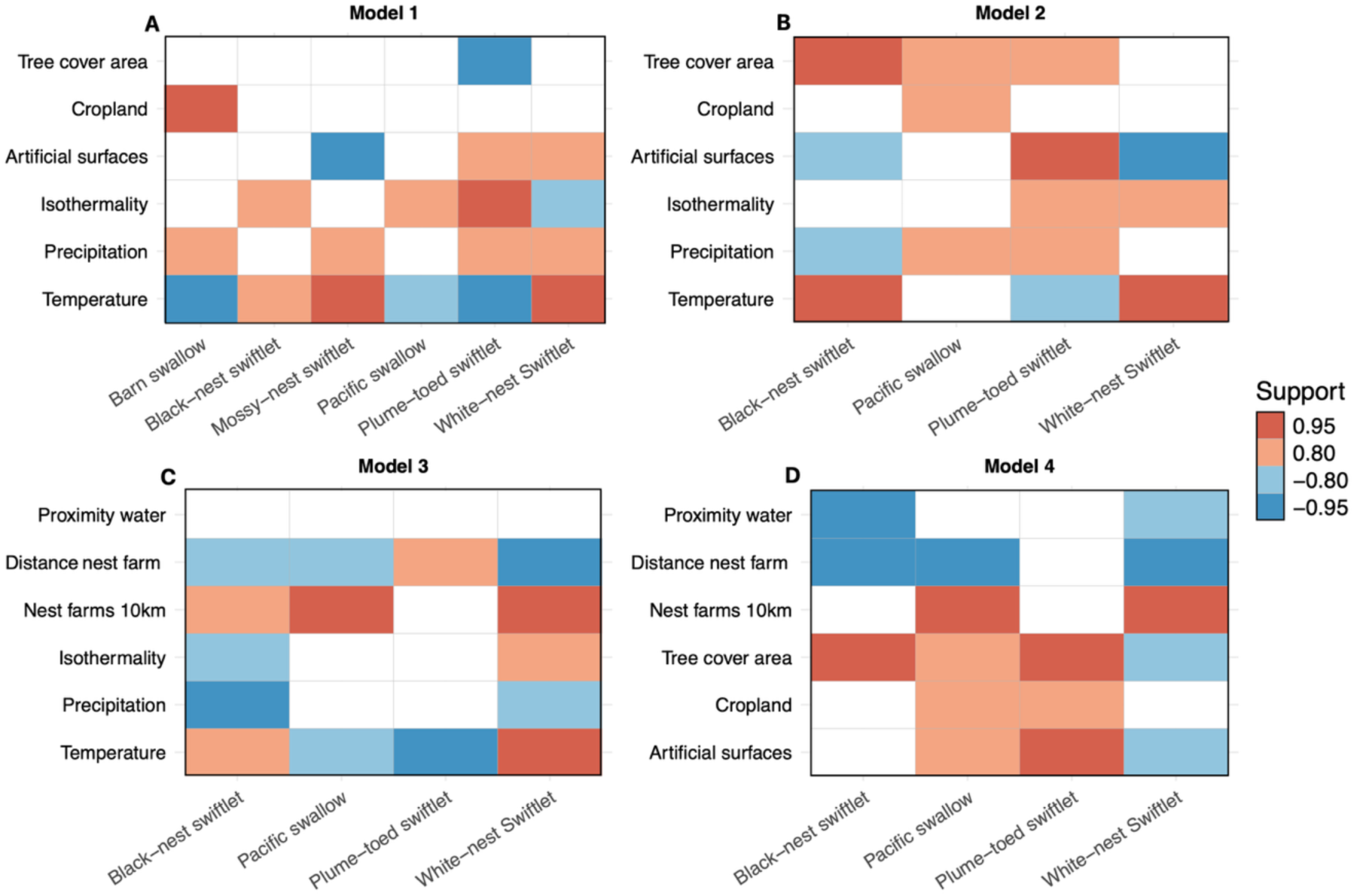
Posterior β estimates of species–environment relationships across four JSDMs. Each panel corresponds to a different model (Models 1-4, A-D). Rows represent environmental predictors, and columns represent species. Cell shading reflects the posterior mean of each β estimate, with credible intervals derived from the posterior distributions of the β coefficients. Red indicates positive associations and blue indicates negative associations. Only associations with moderate (posterior support 0.80–0.95) or strong (posterior support >0.95) credibility are shown. Posterior support reflects the proportion of MCMC samples for which the sign of the effect (positive or negative) was consistent, thus representing certainty of direction of species-environment response.

**Table 2.** Variance partitioning and standardized β parameter estimates for each species in four JSDMs. Variance partitioning shows the proportion of explained variance in species occurrence attributed to each environmental covariate, based on Tjur’s R^2^. Values in parentheses give the posterior mean of the standardized β coefficients. Bolded values with double asterisks (**) indicate strong posterior support for the direction of the effect (posterior probability > 0.95), while single asterisks (*) indicate moderate support (posterior probability 0.80-0.95).

| Model | Species | Temp. | Precip. | Isotherm. | Artificial surfaces | Cropland | Tree cover | Random effect |
| --- | --- | --- | --- | --- | --- | --- | --- | --- |
| Model 1: | Barn | 0.033 | 0.011 | 0.008 | 0.011 | 0.033 | 0.014 | 0.882 |
| Citizen science | swallow | **(-0.007\*\*)** | (0.001*) | (-0.000) | (-0.008) | **(0.016\*\*)** | (0.000) | **(20.50\*\*)** |
|  | Pacific | 0.014 | 0.004 | 0.014 | 0.004 | 0.004 | 0.009 | 0.947 |
|  | swallow | (-0.007*) | (0.000) | (0.003*) | (-0.001) | (0.003) | (0.000) | (18.383*) |
|  | White-nest | 0.150 | 0.035 | 0.018 | 0.021 | 0.008 | 0.013 | 0.742 |
|  | swiftlet | **(0.025\*\*)** | (0.001*) | (-0.002*) | (0.022*) | (-0.003) | (0.001) | (-78.280*) |
|  | Plume-toed | 0.058 | 0.007 | 0.037 | 0.011 | 0.005 | 0.023 | 0.849 |
|  | swiftlet | **(-0.013\*\*)** | (0.000*) | **(0.004\*\*)** | (0.013*) | (0.002) | **(-0.001\*\*)** | **(35.26\*\*)** |
|  | Black-nest | 0.093 | 0.008 | 0.053 | 0.010 | 0.013 | 0.012 | 0.776 |
|  | swiftlet | (0.024*) | (0.000) | (0.007*) | (-0.008) | (0.012) | (0.002) | **(-82.54\*\*)** |
|  | Mossy-nest | 0.116 | 0.024 | 0.014 | 0.030 | 0.008 | 0.016 | 0.777 |
|  | swiftlet | **(0.025\*\*)** | (0.000*) | (0.002) | **(-0.027\*\*)** | (0.001) | (0.000) | **(-82.28\*\*)** |
| Model 2: | Pacific | 0.087 | 0.141 | 0.134 | 0.056 | 0.128 | 0.096 | 0.358 |
| Survey | swallow | (0.157) | (0.268*) | (0.257*) | (0.087) | (0.232*) | (0.191*) | **(0.378\*\*)** |
|  | White-nest | 0.789 | 0.027 | 0.009 | 0.085 | 0.052 | 0.017 | 0.022 |
|  | swiftlet | **(1.270\*\*)** | (-0.202) | (0.036*) | **(-0.375\*\*)** | (0.263) | (-0.102) | (0.154*) |
|  | Plume-toed | 0.142 | 0.275 | 0.134 | 0.154 | 0.046 | 0.138 | 0.111 |
|  | swiftlet | (-0.264*) | (0.434*) | (0.277*) | **(0.309\*\*)** | (-0.075) | (0.276*) | **(0.384\*\*)** |

|  | Black-nest<br>swiftlet | 0.824<br>**(2.379\*\*)** | 0.038<br>**(-0.453\*)** | 0.006<br>**(-0.099)** | 0.046<br>**(-0.485\*)** | 0.010<br>**(0.031)** | 0.067<br>**(0.633\*\*)** | 0.009<br>**(-1.860\*\*)** |
| --- | --- | --- | --- | --- | --- | --- | --- | --- |
|  |  | Temp. | Precip. | Isotherm. | Distance<br>water | Distance<br>nest farm | Nest farm<br>10km | Random<br>effect |
| Model 3: | Pacific | 0.023 | 0.060 | 0.093 | 0.041 | 0.033 | 0.612 | 0.138 |
| Survey | swallow | **(-0.190\*)** | **(0.127)** | **(-0.067)** | **(0.057)** | **(-0.206\*)** | **(0.931\*\*)** | **(0.368\*\*)** |
|  | White-nest<br>swiftlet | 0.196<br>**(0.797\*\*)** | 0.015<br>**(-0.188\*)** | 0.012<br>**(0.153\*)** | 0.012<br>**(0.127)** | 0.221<br>**(-0.874\*\*)** | 0.536<br>**(1.047\*\*)** | 0.008<br>**(0.107)** |
|  | Plume-toed<br>swiftlet | 0.180<br>**(-0.355\*\*)** | 0.210<br>**(0.090)** | 0.089<br>**(0.112)** | 0.258<br>**(-0.106)** | 0.100<br>**(0.171\*)** | 0.033<br>**(0.081)** | 0.130<br>**(0.624\*\*)** |
|  | Black-nest<br>swiftlet | 0.301<br>**(0.943\*)** | 0.133<br>**(-0.664\*\*)** | 0.035<br>**(-0.212\*)** | 0.078<br>**(-0.048)** | 0.355<br>**(-1.327\*)** | 0.070<br>**(0.172\*)** | 0.028<br>**(-2.456\*\*)** |
|  |  | Artificial<br>surfaces | Cropland | Tree<br>cover | Distance<br>water | Distance<br>nest farm | Nest farm<br>10km | Random<br>effect |
| Model 4: | Pacific | 0.053 | 0.066 | 0.026 | 0.018 | 0.048 | 0.671 | 0.117 |
| Survey | swallow | **(0.177\*)** | **(0.175\*)** | **(0.051\*)** | **(0.832\*\*)** | **(-0.021\*\*)** | **(0.175\*\*)** | **(0.402\*\*)** |
|  | White-nest<br>swiftlet | 0.016<br>**(-0.126\*)** | 0.036<br>**(0.235)** | 0.030<br>**(-0.230\*)** | 0.083<br>**(1.537\*)** | 0.314<br>**(-0.445\*\*)** | 0.509<br>**(-0.153\*)** | 0.012<br>**(0.604\*)** |
|  | Plume-toed<br>swiftlet | 0.224<br>**(0.299\*\*)** | 0.070<br>**(-0.114)** | 0.099<br>**(0.195\*\*)** | 0.307<br>**(-0.131\*)** | 0.076<br>**(0.471\*)** | 0.089<br>**(-0.129)** | 0.125<br>**(0.326\*\*)** |
|  | Black-nest<br>swiftlet | 0.138<br>**(-0.195)** | 0.077<br>**(0.012)** | 0.125<br>**(0.199\*\*)** | 0.410<br>**(-0.202\*\*)** | 0.111<br>**(-0.446\*\*)** | 0.048<br>**(0.038)** | 0.090<br>**(-1.264\*\*)** |

### Model 2: Survey-based occurrence records with climate and land-use predictors

Model 2, which replaced citizen-science records with structured survey data, had a lower AUC and Tjur’s R^2^ and higher RMSE for in-sample data than Model 1, but still reflected good discriminatory performance and moderate explanatory power (AUC = 0.880; Tjur’s R^2^ = 0.26, RMSE = 0.277, Table 1). Furthermore, it had much better predictive performance on withheld data (mean Tjur’s R² = 0.193; Table 1), although mean CV AUC remained a bit low (0.631). The spatial random effect in Model 2 accounted for far less variance compared to Model 1, indicating that structured surveys substantially reduced residual spatial correlation (Figure 3B). There was strong posterior support for several species-environment associations with larger β parameter values than Model 1 (Figure 4B, Table 2, Supplemental Material). After accounting for environmental effects, the Ω parameter revealed a moderately supported negative residual correlation (Ω ≈ 0.806) between the Plume-toed swiftlet and Pacific swallow (Supplemental Material). This association suggests that these two species co-occurred less frequently than expected by chance.

### Model 3: Survey-based occurrence records with climate and landscape features

Model 3, which included structured survey data, climate variables, and landscape features, was the best performing model. It had slightly higher RMSE and slightly lower AUC and Tjur’s R^2^ than Model 2 for in-sample data (RMSE = 0.298; AUC =0.861, Tjur’s R^2^ = 0.234; Table 1) but the highest cross-validated metrics (mean AUC = 0.712; mean Tjur’s R² = 0.201; Table 1). This suggests that species-specific landscape features yielded a more accurate model with moderate predictive performance. Furthermore, after including landscape features, the negative residual correlation between Plume-toed swiftlet and Pacific swallow in Model 2 was no longer supported (Supplemental Material). The signal in Model 2 thus likely reflected divergent responses to landscape features, specifically nest farms.

Variance partitioning for Model 3 revealed that the number and proximity of nest farms accounted for substantial variance (Figure 3C, Table 2). For the Pacific swallow, the number of nest farms within 10 km explained 61% of the variance in occurrences, with no other covariate explaining more than 9% (Figure 3C). For the White-nest swiftlet, number and distance to nest farms together explained 76% of the variance, and mean annual temperature contributed 20%. Occurrence of the Black-nest Swiftlet was influenced by distance to nest farms (35.5%), mean annual temperature (30.1%), and precipitation seasonality (13.3%). Only the Plume-toed swiftlet was not strongly influenced by nest farms (13%), with occurrence primarily explained by temperature (18%), distance to water (25.8%), and the spatial random effect (19%).

Inspection of β parameters showed that White-nest swiftlet and Pacific swallow exhibited strong positive responses to the number of nest farms within 10 km (Figure 4C, Table 2). Black-nest and White-nest swiftlets and Pacific swallows also exhibited strong or moderate negative responses to distance from nest farms, indicating they occurred more frequently closer to farms (Figure 4C, Table 2). However, the Plume-toed swiftlet showed a moderate positive response, indicating occurrences declined closer to nest farms (Figure 4C, Table 2). The Plume-toed swiftlet also showed a negative response to mean annual temperature, and the Black-nest swiftlet displayed a negative response to precipitation seasonality.

Because distance to nest farms was strongly or moderately supported for all species, we visualized the effects of this variable on probability of occurrence using partial dependence plots (Figure 5). Black-nest swiftlets exhibited low mean predicted occurrence near farms, with probability declining with distance (Figure 5A), consistent with the rarity of this species in our surveys. Plume-toed swiftlets displayed the opposite pattern, with higher predicted occurrence farther from farms (Figure 5B), suggesting potential avoidance behavior. White-nest swiftlets and Pacific swallows had high probability of occurrence near farms that declined with distance, with a steeper drop for White-nest swiftlets (Figure 5C) than for Pacific swallows (Figure 5D).

**Figure 5.**
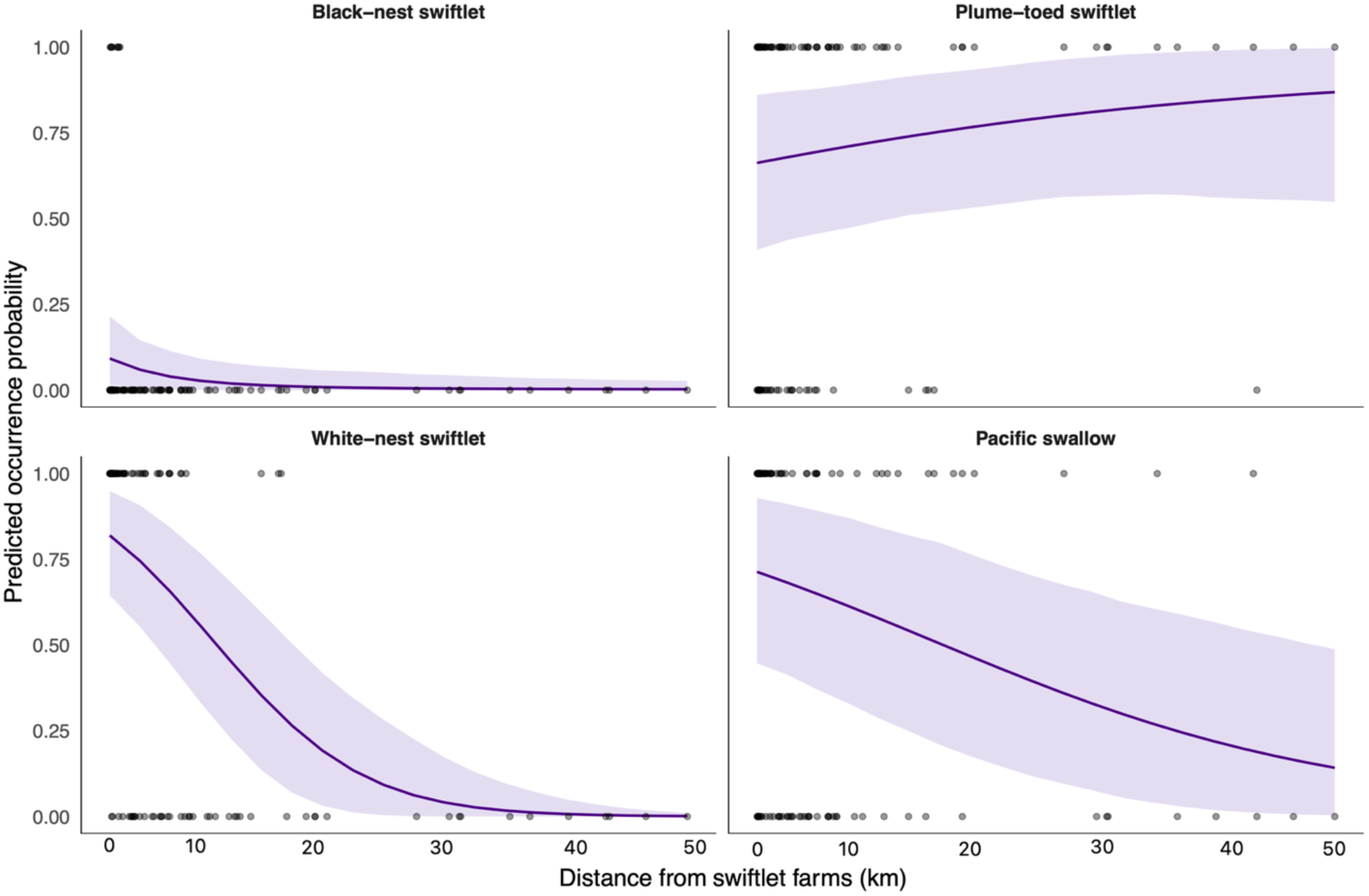
Partial dependence plots for each species estimated from Model 3. Plots show predicted occurrence probabilities (solid line) with 95% confidence intervals (purple shading) for the four focal species as a function of distance from nest farms in the best-fit model (Model 3). Points show observed occurrence values. The White-nest and Black-nest swiftlets (A,C) and Pacific swallow (D) exhibit higher occurrence probabilities near farms, whereas the Plume-toed swiftlet (C) shows decreasing probabilities with closer proximity. These contrasting responses highlight divergent habitat preferences and suggest potential shifts in coexistence dynamics within farm-dense landscapes.

### Model 4: Survey-based occurrence records with land-use and landscape features

Compared to Model 3, Model 4 retained species-specific landscape features but replaced climate predictors with land-use data. While in-sample performance matched that of Model 3, Model 4 showed slightly lower predictive performance on withheld data (mean Tjur’s R^2^ = 0.160, mean AUC = 0.687; Table 1). Landscape features accounted for similar or greater proportions of variance and species-environment relationships had similar posterior support compared to Model 3 for most species (Figure 3D, Figure 4D, Table 2). For the Plume-toed swiftlet, artificial surfaces explained 22.4% of variance in occurrence (Figure 3D, Table 2) and the proportion of variance explained by the random effect declined relative to Model 3, suggesting land cover is important for predicting occurrence of this species. After accounting for environmental factors, the Ω parameter revealed no strong residual correlations between species, consistent with Model 3 (Supplemental Material).

### Inclusion of morphological, behavioral, and phylogenetic data in JSDMs

Integrating morphological and behavioral traits into JSDMs improved in-sample measures of model performance but resulted in slightly lower performance on withheld data (Table 1). The strength of morphological trait–environment associations was generally moderate across models, with few consistent trends (Supplemental Material). When behavioral traits related to foraging height were incorporated into Models 2–4 in place of morphological traits, all trait–environment associations remained moderate (Supplemental Material). Minimum foraging height was consistently negatively associated with artificial surface cover across Models 2 and 4, while maximum foraging height showed a positive association, suggesting that species foraging at higher strata use urban environments less frequently. Both trait sets exhibited low median values for phylogenetic signal (0.02 for morphology, 0.00 for behavior).

## Discussion

In this study, we used multiple modeling approaches to evaluate how human land-use has shaped interspecific co-occurrence in an aerial insectivore community in Southeast Asia. We found high geographic niche overlap among species that exploit anthropogenic environments, which is evidence that human land-use change may intensify competition among ecologically similar species. We tested this hypothesis using a series of JSDMs that allowed us to partition shared environmental responses and isolate residual co-occurrence correlations, providing deeper insight into potential biotic interactions. The best-performing model combined climate variables with field-measured landscape features (number and distance to nest farms) and had no strongly supported residual correlations. Our models thus show that anthropogenic activity, particularly the creation of artificial nesting sites, broadly shapes patterns of co-occurrence across our study region, but appears to act through environmental filtering rather than direct competition or facilitation. Below, we discuss the implications of these findings, the potential for land-use change to restructure community dynamics, and the limitations and conservation applications of our modeling approach.

### Geographic niche overlap

Human activity has dramatically increased available nest sites for swiftlets and swallows in Southeast Asia. If this has induced more frequent interspecific co-occurrence, we hypothesized that we would find higher predicted geographic niche overlap in SDMs that included land-use predictors compared to those that did not, particularly for species that exploit anthropogenic environments. We indeed found this to be the case, with Pacific swallows, Barn swallows, and Plume-toed swiftlets, the species that most regularly use human-modified environments, exhibiting particularly high niche overlap. However, the best-fit SDMs for each species included both land-use and climate predictors, which suggests that environmental filtering related to abiotic preferences persists among species and may mitigate competition to some extent. Previous studies have used geographic niche overlap to infer processes influencing species coexistence. For example, comparisons of the known ranges of Palearctic bat species with those predicted by SDMs found that two ecologically similar pairs of cryptic species showed spatial segregation consistent with competitive exclusion (Novella-Fernandez et al. 2021). Furthermore, reproductive interference has been proposed to limit niche space in sympatric Rubyspot damselflies (*Hetaerina spp.*, Grether et al., 2024). By examining geographic niche overlap under different environmental predictor sets, our results suggest that continued land-use change could further homogenize niches and increae the frequency of competitive interactions.

### Effects of citizen-science records vs. structured surveys on JSDM predictions

Unstructured and semi-structured citizen-science data offer a unique opportunity to model the distributions of species that are wide-ranging or difficult to detect. However, applications in JSDMs have been largely limited to well-structured programs like the French and Spanish Breeding Bird Surveys (Planillo et al. 2021; Zurell 2020). Because the models we used to calculate geographic niche overlap used citizen-science records from eBird, we tested whether JSDMs using the same unstructured occurrence records could perform comparably to those using structured survey data. To our knowledge, ours is among the first studies to explore this application of eBird data.

Our results showed that models using eBird records were overfit, exhibited strong spatial random effects, and generalized poorly relative to models using structured survey data. We suspect this poor performance stems from three related factors. First, citizen-science data often have strong spatial sampling bias, with observations clustering around human-populated areas (Romera-Romera & Nieto-Lugilde 2025). Second, eBird occurrences in our focal region were patchily distributed, exacerbating sampling bias. Third, species like the Plume-toed swiftlet and Pacific swallow thrive in human-modified environments, meaning their distributions can be further confounded by spatial bias in sampling effort (Ramirez et al. 2025). While single-species SDMs can control for spatial sampling bias through, for example, choice of background selection methods (Merow et al., 2013; Barbet-Massin et al. 2012; Ramirez et al. 2025), controlling for bias is more difficult with community-level data (D’Amen et al. 2015; Buckland & Johnston 2017). Future use of citizen-science data in JSDMs should focus on regions with comprehensive sampling coverage for the focal species to help mitigate these biases.

### Effects of field-measured landscape features on model predictions

JSDMs focused on large spatial scales typically use relatively general environmental predictors such as climate and land cover, which can capture coarse niche differences (e.g., Cranston et al., 2022; Antão, Weigel, Strona et al., 2022). At finer spatial scales, where biotic interactions are expected to be more important to community assembly, it is more common to incorporate species-specific or microhabitat characteristics to better separate biotic interactions from shared environmental responses (Wagner et al., 2020; Elo et al., 2021). Although the scale of our study was somewhat small compared to the ranges of the focal species, it nonetheless covered a large spatial extent, reflective of the high dispersal ability of swiftlets and swallows. We therefore compared the performance of models that used general, satellite-derived predictors to those that used species-specific landscape features measured in the field.

We found that while models fit with only climate and land-use variables performed moderately well, including the number and proximity of nest farms enhanced predictive power and reduced residual correlations between species (Model 3). Indeed, these features explained the most variance in occurrences. White-nest swiftlets, Black-nest swiftlets, and Pacific swallows all had higher predicted occurrence near nest farms. In contrast, Plume-toed swiftlets were negatively associated with nest farms, indicating potential avoidance. Crucially, the models including landscape features eliminated the negative residual co-occurrence between Plume-toed swiftlets and Pacific swallows in Model 2, indicating that this association was likely driven by divergent responses to nest farms rather than direct competitive interactions.

Although nest farms explained the most variance in occurrences, different responses to climate and land-use variables in Models 2, 3, and 4 suggest divergence in habitat preferences that may facilitate coexistence. Both White-nest and Black-nest swiftlets were positively associated with temperature and negatively associated with artificial surfaces but differed in their responses to other variables. Most notably, Black-nest swiftlets were positively associated with tree cover, consistent with more common use of caves compared to White-nest swiftlets, which were more often near nest farms. In contrast to the two farmed species, Pacific swallows and Plume-toed swiftlets exhibited negative associations with temperature and positive associations with all land cover types, including artificial surfaces and cropland. This likely reflects the frequent use of structures such as bridges, culverts, and buildings as nest sites.

Overall, our models had moderate explanatory power and showed that access to artificial nesting structures is a pivotal driver of occurrence in swiftlets and swallows, potentially reshaping their realized niches. By providing abundant nesting opportunities, nest farms likely boost local swiftlet population densities and allow access to previously unavailable habitat, which may in turn intensify competition for shared insect prey and could ultimately lead to declines in competitor species. Although our study does not include abundance data, abundance likely plays an important role in mediating species interactions. We observed dramatic variation in abundance of the different species, with some surveys recording hundreds of individuals.

However, due to high overdispersion across sites and the challenges of accurately recording abundance in large mixed-species flocks, integrating such data into JSDMs remains difficult.

### Inclusion of morphology, behavior, and phylogeny in JSDMs

We incorporated morphological and behavioral traits and a phylogenetic tree into our models to provide further insight into drivers of co-occurrence. Phylogenetic signal was low across models, suggesting that trait-based niche variation reflects functional convergence rather than conserved responses. Incorporating traits into JSDMs enhanced explanatory performance for in-sample data, likely because traits further explained species–environment and species–species relationships among ecologically similar taxa. However, predictive performance on withheld data declined slightly. This likely reflects a combination of increased model complexity and weak or context-dependent trait-environment relationships. This is particularly likely for behavioral traits, which are expected to vary with local conditions– a pattern not captured by JSDMs.

### Biotic interactions and environmental filtering

We predicted that if human land-use change is altering biotic interactions among swiftlets and swallows, we would observe negative residual associations reflective of competition after accounting for shared responses to environmental predictors. However, our environmental predictors explained substantial proportions of variance in occurrences without strongly supported residual correlations. This does not mean that human activity is not affecting these communities; rather, it indicates that the landscape features included in our models sufficiently account for variation in occurrence. Indeed, we find that nest farms explained most of this variance, underscoring how dramatically human activity has shaped the distribution of these species.

The absence of residual correlations also does not imply a lack of competition; instead, it suggests that competitive exclusion may already be embedded within the species-environment relationships. For instance, Plume-toed swiftlets may avoid competition with larger swiftlets and swallows not through direct interactions, but by occupying urban environments farther from nest farms. JSDMs are not intended to provide definitive evidence of species interactions (D’Amen 2018; Wilkinson 2020): residual correlations, whether positive or negative, represent potential associations, which may reflect shared microhabitat preferences, missing covariates, or true biotic interactions. Similar findings have been reported in other systems, such as tropical frog communities, where positive residuals were due to shared microhabitat use rather than facilitation (Pollock et al. 2014), and in pipits, where negative residual correlations were attributed to indirect competition (Bastianelli et al. 2017).

### Conservation implications

Our results show that anthropogenic nesting sites strongly influence the composition of aerial insectivore communities in Borneo. This has important implications for the conservation and management of these species, particularly the economically important swiftlets. Prior to the boom in nest farming, cave-based populations of White-nest and Black-nest swiftlets were declining due to overharvesting (Medway 1963, Lim & Cranbrook 2002). These declines were significant: at Niah, Malaysia, Black-nest swiftlets were reduced from over 3 million birds to just 300,000 in 50 years (Gausset 2004; Stimpson 2013), and many smaller cave-based colonies were extirpated entirely (Lim & Cranbrook 2002). Commercial nest farming began in the 1990s and has exploded throughout Mainland and Insular Southeast Asia, with more than 60,000 farms estimated in Malaysia alone (Malaysia Economic Transformation Programme Annual Report, 2013). Nest farms have likely helped slow declines driven by cave overharvesting and have been touted by some as a form of sustainable development (Ito et al., 2021); the nest trade now generates billions of dollars per year in economic activity across Southeast Asia (Thorburn 2015). However, it is unclear whether swiftlet populations have rebounded to their historic levels, or if they remain below carrying capacity. Furthermore, much remains unknown about the consequences of nest farming for the broader avian community, including how deforestation and the expansion of nest farming jointly influence foraging ranges and dietary overlap, whether shifts in the distribution and local abundance of farmed swiftlets exacerbate competition with local non-farmed species, and whether farmed and cave populations interbreed or compete with each other. Here we use occurrence data to present the first study of the impacts of nest farms on aerial insectivore communities. Future work should incorporate complementary analyses, such as diet analysis and abundance surveys. This would further determine whether the patterns of co-occurrence we report here reflect persistent habitat filtering or emerging competition, and would help inform efforts to sustainably manage swiftlet populations.

## Acknowledgements

We are grateful to the Danau Girang Field Centre, KOPEL, and the Sabah Wildlife Department for support in the field. Research was conducted under Sabah Biodiversity Centre Access License numbers JKM/MBS.1000-2/2 JLD.19(8) and JKM/MBS.1000-2/2/1 JLD.1(130). A. Hernandez and I. Garcia assisted with classification of landcover. A. Hund, M. Tingley and R. Blakey provided valuable feedback throughout the project. Funding was provided by NSF-DEB 1947306 to E. Scordato.

