## Supplemental material for "Human land-use change drives co-occurrence of ecologically similar avian aerial insectivores in Southeast Asia"

### Supplementary Methods

*Citizen science occurrence data*

We downloaded records from the eBird database (Sullivan et al., 2009) from 2015 to 2024 for each of our six focal species. To control for sampling bias, we used the R package *auk* (Strimas-Mackey et al., 2018) to filter occurrence data according to best practices (Johnston et al., 2021). Specifically, data were restricted to surveys conducted via traveling or stationary count protocols, with traveling distances of 5 km or less, durations of 300 minutes or fewer, and group sizes of no more than 10 observers. Only complete checklists (i.e., those that reported all species detected during surveys) were retained.

*SDM Construction and Evaluation*

We built SDMs for each focal species using eBird occurrence records. We thinned the occurrence records so that only one record was retained per 1 $km^2$ raster cell. Background points were sampled from every grid cell in the study region (n = 20,000; Table S1), a method shown to have superior performance for human-associated species (Ramirez et al. 2025).

Following Ramirez et al. (2025), we built three sets of SDMs for each species reflecting different land-use scenarios: models that included only climate variables from CHELSA ("climate-only"), models with only land use variables from the GLC ("land-use"), and models with all variables ("both sets"). SDMs were generated using Maxent Java software (v3.4.1; Phillips et al., 2017) implemented via the ENMeval package (v2.0.4; Kass et al., 2021), yielding a total of 18 models across our focal species (3 models each for six species). Model parameters were tuned using regularization multipliers ranging from 0-5 at 0.5-point increments and feature class combinations of 'Linear', 'Quadratic', 'Hinge', 'Linear + Quadratic', 'Quadratic + Hinge' and 'Linear + Quadratic + Hinge'. We employed the "block" method for spatial cross-validation (Muscarella et al., 2014).

Within each predictor set, we selected the optimal model for each species by identifying the feature class and regularization multiplier combination that had the highest value of the Continuous Boyce Index (CBI), which ranges from –1 to 1 and reflects model calibration quality (Hirzel et al., 2006). For each of these best models, we further assessed model performance by generating 500 null models using the ENMnulls function (Raes & ter Steege, 2007; Bohl et al., 2019). Empirical models exceeding the 95$^{th}$ quantile of the null distribution CBI distribution were considered a good fit. Null modeling showed that all SDMs exceeded the 95% quantile of the null distribution for CBI, indicating good model fits. The only exception was the climate-only model for the Plume-toed swiftlet, for which the empirical model only exceeded the 80% quantile. Finally, for each species we chose the best-performing variable set (climate only, land-use only, or both sets) using AICc.

### Supplementary Results

Below we present interpretations of the beta parameters for Models 1, 2, and 4. In HMSC, support for parameter estimates is measured by the proportion of posterior samples above or below zero, and thus even small mean estimates of the parameters can be strongly supported if variance is low and their 95% credible intervals do not overlap zero.

#### *Model 1*

Amongst environmental predictors, mean annual temperature explained the greatest proportion of variance in species occurrence and was strongly negatively associated with occurrence for

Barn swallow, Pacific swallow, and Plume-toed swiftlet, and strongly positively associated for Black-nest, Mossy-nest, and White-nest swiftlets; however, values for the parameter estimates were small (Figure 4A, Table 2).

*Model 2*

The Black-nest swiftlet displayed a strong positive association with mean annual temperature and tree cover, and moderately negative responses to precipitation seasonality and artificial surfaces (Figure 4B, Table 2). Similarly, the White-nest swiftlet had a strong positive response to temperature and a strong negative response to artificial surfaces. By contrast, Plume-toed swiftlet**s** were moderately negatively associated with temperature and positively associated with artificial surfaces and cropland. Pacific swallow**s** were moderately positively influenced by precipitation seasonality, cropland and tree cover.

*Model 4*

Black-nest swiftlets showed a strong positive response to tree cover, and strong negative relationships with proximity to water and nest farms; these relationships were stronger than in Model 3 (Figure 4D, Table 2). White-nest swiftlets showed similar responses to nest farms in Model 4 as in Model 3 (Figure 4D, Table 2). Pacific swallows also showed similar responses to nest farms in Models 3 and 4, plus a moderately positive association with artificial surfaces that was absent from Model 2. Finally, Plume-toed swiftlets displayed strong positive responses to both artificial surfaces and tree cover in Model 4, consistent with Model 2 (Figure 4D, Table 2), as well as a new, moderately positive relationship with cropland. The moderate positive response with increasing distance from nest farms in Model 3 disappeared in Model 4.

**Table S1**. *Species occurrences used in different modeling approaches.* SDMs used presence data derived from eBird records, with 20,000 randomly sampled background points included per species. JSDMs using citizen science data used a community presence-absence matrix that included presence records and zero-filled absences from all complete eBird checklists in our study region (n = 1,430). JSDMs using structured survey data included 155 field surveys, where absence represents surveyed locations where species were not detected.

| **Model** | **Count Type** | **Barn swallow** | **Pacific swallow** | **White-nest swiftlet** | **Plume-toed swiftlet** | **Black-nest swiftlet** | **Mossy-nest swiftlet** |
|---|---|---|---|---|---|---|---|
| SDM | Presence | 146 | 312 | 51 | 266 | 50 | 32 |
| | Absence | 20,000 | 20,000 | 20,000 | 20,000 | 20,000 | 20,000 |
| JSDM (Citizen science) | Presence | 146 | 312 | 51 | 266 | 50 | 32 |
| | Absence | 1021 | 855 | 1116 | 901 | 1117 | 1135 |
| JSDM (Structured survey) | Presence | N/A | 92 | 93 | 95 | 20 | 3 |
| | Absence | N/A | 63 | 62 | 60 | 135 | 152 |

**Table S2.** *Summary of environmental and trait covariates included across the five JSDMs*. Model specifications differ by the data source and predictor sets: Model 1 (M1) used citizen-science occurrences with climate and GLC land-cover predictors; M2 used structured survey occurrences and climate and land cover variables; M3 used structured survey occurrences, climate variables, and landscape features; and M4 used structured survey occurrences, land-use variables, and landscape features. "X" indicates inclusion of the variable in the model.

| Variable | (M1) Citizen Science ~ Climate + Land-use | (M2) Survey occurrences ~ Climate + Land-use | (M3) Survey occurrences ~ Climate + Landscape features | (M4) Survey occurrences ~ Land-use + Landscape features |
|---|---|---|---|---|
| Mean annual temperature | X | X | X | |
| Mean annual precipitation | X | X | X | |
| Isothermality | X | X | X | |
| Artificial surfaces (prop. cover) | X | X | | X |
| Cropland (prop. cover) | X | X | | X |
| Tree cover (prop. cover) | X | X | | X |
| Proximity to water (km) | | | X | X |
| Proximity to nearest nest farm (km) | | | X | X |
| Number of nest farms within $10km^2$ grid cells | | | X | X |
| Behavioral traits (Minimum, maximum and modal foraging strata) | | X | X | X |
| Morphometric traits (Beak culmen, mass, wing length, beak depth) | X | X | X | X |

**Table S3.** *Optimal SDM parameters and performance metrics for each focal species across predictor sets.* Optimal models were selected by identifying the tuning combination that maximized the empirical Continuous Boyce Index (CBI). For each retained model, we report the selected feature class (FC), regularization multiplier (rm), Area Under the Curve (AUC), and empirical CBI. Higher AUC and CBI values indicate stronger discrimination and calibration performance, respectively.

| Species | Predictor Set | FC | RM | AUC | CBI | AICc | ΔAICc |
|---|---|---|---|---|---|---|---|
| Barn swallow | Both predictor sets | H | 2 | 0.78 | 0.83 | 2512.45 | 50.66 |
| | Climate only | LQHPT | 0.5 | 0.74 | 0.77 | 3852.14 | 1390.35 |
| | **Land-use only** | **LQHP** | **0.5** | **0.71** | **0.83** | **2461.79** | **0.00** |
| Pacific swallow | **Both predictor sets** | **LQP** | **1** | **0.79** | **0.84** | **3274.74** | **0.00** |
| | Climate only | H | 0.5 | 0.74 | 0.80 | 3385.35 | 110.61 |
| | Land-use only | LQP | 1.5 | 0.74 | 0.84 | 3357.98 | 83.24 |
| White-nest swiftlet | **Both predictor sets** | **L** | **3.5** | **0.80** | **0.86** | **925.88** | **0.00** |
| | Climate only | H | 2.5 | 0.81 | 0.81 | 933.84 | 7.96 |
| | Land-use only | LQP | 1 | 0.76 | 0.82 | 952.88 | 26.99 |
| Plume-toed swiftlet | **Both predictor sets** | **H** | **1** | **0.80** | **0.86** | **2603.45** | **0.00** |
| | Climate only | LQHP | 2.5 | 0.76 | 0.81 | 2625.52 | 22.07 |
| | Land-use only | LQHP | 0.5 | 0.72 | 0.82 | 2651.98 | 48.53 |
| Black-nest swiftlet | **Both predictor set** | **L** | **4** | **0.73** | **0.73** | **780.32** | **0.00** |
| | Climate only | LQHPT | 0.5 | 0.69 | 0.74 | 788.96 | 4.42 |
| | Land-use only | LQHPT | 1 | 0.66 | 0.67 | 784.74 | 8.64 |
| Mossy-nest swiftlet | **Both predictor sets** | **L** | **0.5** | **0.75** | **0.81** | **597.38** | **0.00** |
| | Climate only | H | 1.5 | 0.75 | 0.73 | 599.54 | 2.16 |
| | Land-use only | LQ | 0.5 | 0.72 | 0.75 | 601.36 | 3.98 |

**Table S4.** *In-sample summary statistics for each species in each model.*

| Model | Statistic | White-nest swiftlet | Black-nest swiftlet | Plume-toed swiftlet | Pacific swallow | Mossy-nest swiftlet | Barn swallow |
|---|---|---|---|---|---|---|---|
| Model 1 Citizen science | RMSE | 0.196 | 0.171 | 0.324 | 0.283 | 0.159 | 0.343 |
| | AUC | 0.976 | 0.979 | 0.938 | 0.981 | 0.986 | 0.895 |
| | Tjurs R2 | 0.516 | 0.497 | 0.472 | 0.558 | 0.281 | 0.345 |
| Model 2 Structured survey | RMSE | 0.259 | 0.222 | 0.347 | 0.279 | N/A | N/A |
| | AUC | 0.894 | 0.802 | 0.853 | 0.970 | N/A | N/A |
| | Tjurs R2 | 0.236 | 0.220 | 0.265 | 0.330 | N/A | N/A |
| Model 3 Structured survey | RMSE | 0.291 | 0.265 | 0.366 | 0.270 | N/A | N/A |
| | AUC | 0.890 | 0.881 | 0.801 | 0.873 | N/A | N/A |
| | Tjurs R2 | 0.335 | 0.138 | 0.222 | 0.242 | N/A | N/A |
| Model 4 Structured survey | RMSE | 0.303 | 0.181 | 0.370 | 0.381 | N/A | N/A |
| | AUC | 0.900 | 0.899 | 0.760 | 0.813 | N/A | N/A |
| | Tjurs R2 | 0.380 | 0.192 | 0.153 | 0.213 | N/A | N/A |

**Table S5.** *Pairwise comparisons of geographic overlap: both variable sets.* Grey boxes show geographic overlap estimates of SDMs based on Schoener's D statistic. These are the best-fit models for all species, and show fairly high geographic niche overlap among all species.

| Species | Barn swallow | Pacific swallow | White-nest swiftlet | Plume-toed swiftlet | Black-nest swiftlet | Mossy-nest swiftlet |
|---|---|---|---|---|---|---|
| Pacific swallow | 0.77 | | | | | |
| White-nest swiftlet | 0.84 | 0.76 | | | | |
| Plume-toed swiftlet | 0.83 | 0.87 | 0.76 | | | |
| Black-nest swiftlet | 0.82 | 0.75 | 0.82 | 0.74 | | |
| Mossy-nest swiftlet | 0.81 | 0.78 | 0.81 | 0.78 | 0.80 | |

**Table S6.** *Pairwise comparisons of geographic overlap: climate only.* Grey boxes show geographic overlap estimates of SDMs based on Schoener's D statistic. Geographic niche overlap among species in models that only include climate data is moderate to high.

| Species | Barn swallow | Pacific swallow | White-nest swiftlet | Plume-toed swiftlet | Black-nest swiftlet | Mossy-nest swiftlet |
|---|---|---|---|---|---|---|
| Barn swallow | | | | | | |
| Pacific swallow | 0.80 | | | | | |
| White-nest swiftlet | 0.68 | 0.77 | | | | |
| Plume-toed swiftlet | 0.76 | 0.89 | 0.79 | | | |
| Black-nest swiftlet | 0.72 | 0.83 | 0.76 | 0.81 | | |
| Mossy-nest swiftlet | 0.67 | 0.79 | 0.85 | 0.83 | 0.81 | |

**Table S7.** *Pairwise comparisons of geographic overlap: human land use only.* Grey boxes show geographic overlap estimates of SDMs based on Schoener's D statistic. Geographic niche overlap among species in models that only include human land-use data is uniformly high, particularly among species that regularly use human-modified environments (Pacific swallow, Plume-toed swiftlet, and Barn swallow).

| Species | Barn swallow | Pacific swallow | White-nest swiftlet | Plume-toed swiftlet | Black-nest swiftlet | Mossy-nest swiftlet |
|---|---|---|---|---|---|---|
| Pacific swallow | 0.88 | | | | | |
| White-nest swiftlet | 0.82 | 0.87 | | | | |
| Plume-toed swiftlet | 0.87 | 0.93 | 0.86 | | | |
| Black-nest swiftlet | 0.82 | 0.87 | 0.86 | 0.86 | | |
| Mossy-nest swiftlet | 0.78 | 0.85 | 0.81 | 0.83 | 0.82 | |

**Table S8.** *Residual Association (Ω) Matrix for the JSDM Model 1-4*. Values represent the mean residual correlations between species. Support values are given in parentheses; support values >0.8 are marked with an asterisk. Mossy-nest and Barn swallow were not included Models 2-4. In model 1, large amounts of unexplained residual variance in the random effect contribute to very large correlations in the residuals.

| Species | Model 1 | Model 2 | Model 3 | Model 4 |
|---|---|---|---|---|
| Barn swallow – Pacific swallow | 0.999 (1.000) | N/A | N/A | N/A |
| Barn swallow- White-nest swiftlet | 0.999 (1.000) | N/A | N/A | N/A |
| Barn swallow -Plume-toed swiftlet | 0.999 (1.000) | N/A | N/A | N/A |
| Barn swallow -Black-nest swiftlet | 0.999 (1.000) | N/A | N/A | N/A |
| Barn swallow - Mossy-nest swiftlet | 0.998 (1.000) | N/A | N/A | N/A |
| Mossy-nest swiftlet – Pacific swallow | 0.999 (1.000) | N/A | N/A | N/A |
| Mossy-nest swiftlet – White-nest swiftlet | 0.999 (1.000) | N/A | N/A | N/A |
| Mossy-nest swiftlet – Plume-toed swiftlet | 0.999 (1.000) | N/A | N/A | N/A |
| Mossy-nest swiftlet – Black-nest swiftlet | 0.999 (1.000) | N/A | N/A | N/A |
| Pacific swallow – White-nest swiftlet | 0.999 (1.000) | 0.174 (0.769) | 0.070 (0.662) | 0.018 (0.589) |
| Pacific swallow – Plume-toed swiftlet | 0.999 (1.000) | -0.256 (**0.806*)** | 0.173 (0.607) | -0.111 (0.550) |
| Pacific swallow – Black-nest swiftlet | 0.999 (1.000) | -0.001 (0.502) | -0.071 (0.467) | -0.081 (0.556) |
| White-nest swiftlet – Plume-toed swiftlet | 0.999 (1.000) | -0.174 (0.410) | -0.118 (0.642) | -0.097 (0.561) |

| | | | | |
|---|---|---|---|---|
| White-nest swiftlet – Black-nest swiftlet | 0.999 (1.000) | 0.002 (0.497) | -0.037 (0.475) | 0.013 (0.506) |
| Plume-toed swiftlet- Black-nest swiftlet | 0.999 (1.000) | 0.007 (0.512) | 0.002 (0.502) | -0.030 (0.512) |

**Table S9.** *Γ Estimates for environmental predictor- morphological trait relationships in models 1-4.* Rows represent environmental predictors, and columns represent morphological trait coefficients. Values reflect the posterior mean estimates. Boldfaced and starred values have support >0.8.

| Model | Predictor | Intercept | Beak Culmen | Mass | Wing Length | Beak Depth |
|---|---|---|---|---|---|---|
| Model 1 | Intercept | **-1.790**** | **1.255*** | -0.537 | **-0.666*** | -0.121 |
| | Temperature | 0.263* | -0.563* | 0.466 | 0.182 | -0.037 |
| | Precipitation | 0.132 | -0.137 | 0.078 | -0.011 | -0.081 |
| | Isothermality | **0.210*** | 0.224 | -0.309 | 0.364 | 0.169 |
| | Artificial surfaces | -0.021 | 0.177 | 0.125 | 0.029 | -0.341 |
| | Cropland | 0.072 | 0.127 | -0.117 | 0.145 | -0.011 |
| | Tree cover area | -0.088 | -0.244 | 0.112 | -0.140 | 0.096 |
| Model 2 | Intercept | **-0.629*** | **1.152*** | 0.074 | **-1.019*** | -0.621 |
| | Temperature | **0.792**** | -0.425 | 0.431 | **0.781*** | -0.137 |
| | Precipitation | 0.149 | 0.306 | 0.140 | **-0.657*** | 0.227 |
| | Isothermality | 0.168 | -0.207 | 0.189 | -0.240 | 0.043 |
| | Artificial surfaces | -0.115 | -0.211 | -0.084 | -0.001 | -0.283 |
| | Cropland | 0.151 | -0.507 | 0.245 | 0.123 | -0.188 |
| | Tree cover area | 0.085 | -0.098 | -0.353 | 0.200 | -0.053 |
| Model 3 | Intercept | -0.133 | 0.428 | 0.589 | -0.215 | -0.543 |
| | Temperature | 0.256 | **-0.503*** | -0.354 | -0.234 | -0.254 |
| | Precipitation | -0.090 | 0.292 | 0.194 | -0.037 | -0.184 |
| | Isothermality | -0.024 | -0.146 | 0.054 | 0.019 | -0.132 |
| | Nest farms 10km | **0.872**** | **0.654*** | -0.119 | -0.521 | -0.062 |
| | Proximity water | 0.064 | 0.028 | -0.033 | -0.152 | -0.092 |
| | Proximity nest farm | **-0.560*** | 0.085 | 0.290 | 0.237 | 0.110 |
| Model 4 | Intercept | **-0.603*** | **1.053*** | -0.077 | **-0.969*** | -0.566 |
| | Artificial surfaces | -0.029 | -0.269 | -0.087 | 0.063 | -0.265 |

| | | | | | |
|---|---|---|---|---|---|
| Cropland | -0.023 | -0.679 | 0.251 | 0.235 | -0.204 |
| Tree cover area | 0.023 | -0.052 | -0.395 | 0.332 | -0.141 |
| Nest farms 10km | **0.609*** | -0.075 | 0.550 | -0.687 | 0.131 |
| Proximity water | **-0.301*** | 0.107 | -0.141 | 0.115 | -0.071 |
| Proximity nest farm | **-1.008**** | 0.066 | -0.174 | **-0.561*** | -0.107 |

**Table S10**. *Γ Estimates for environmental predictor- behavioral trait relationships in models 1-4*. Rows represent environmental predictors, and columns represent behavior trait coefficients. Values reflect the posterior mean estimates. Boldfaced and starred values have support >0.8.

| Model | Predictor | Intercept | Minimum foraging | Max foraging | Modal forgaing |
|---|---|---|---|---|---|
| Model 2 | Intercept | -0.223 | **-0.988**** | 0.528 | -0.401 |
| | Temperature | 0.246 | 0.110 | **0.147*** | 0.482 |
| | Precipitation | -0.121 | -0.268 | -0.004 | -0.190 |
| | Isothermality | -0.029 | -0.059 | 0.168 | -0.021 |
| | Artificial surfaces | **-0.211*** | **-0.054*** | **0.054*** | 0.005 |
| | Cropland | **-0.152*** | -0.004 | 0.029 | 0.019 |
| | Tree cover area | -0.143 | 0.021 | 0.037 | -0.002 |
| Model 3 | Intercept | -0.237 | **-1.236**** | 0.571 | -0.410 |
| | Temperature | 0.245 | 0.123 | **0.145*** | 0.493 |
| | Precipitation | -0.116 | -0.270 | -0.006 | -0.195 |
| | Isothermality | -0.020 | -0.092 | 0.171 | -0.022 |
| | Nest farms 10km | 0.611 | **0.713*** | -0.046 | 0.236 |
| | Proximity water | 0.013 | -0.148 | 0.087 | 0.054 |
| | Proximity nest farm | **-0.547*** | **-0.345*** | **0.265*** | -0.392 |
| Model 4 | Intercept | 3.442 | -1.307 | 0.031 | -1.529 |
| | Artificial surfaces | **-0.200*** | **-0.054*** | **0.055*** | 0.005 |
| | Cropland | **-0.141*** | -0.004 | 0.030 | 0.019 |
| | Tree cover area | -0.143 | 0.021 | 0.039 | -0.003 |
| | Nest farms 10km | -0.090 | 0.011 | 0.032 | -0.022 |
| | Proximity water | -0.271 | **-0.082*** | **0.088*** | -0.037 |
| | Proximity nest farm | 0.202 | -0.091 | -0.058 | 0.048 |

**Figure S1.** *Phylogenetic tree used in JSDMs*. We constructed a maximum likelihood consensus tree from a posterior distribution of 1,000 avian phylogenies obtained from VertLife for our focal species: Barn swallow (*Hirundo rustica*), Pacific swallow (*Hirundo tahitica*), White-nest swiftlet (*Aerodramus fuciphaga*), Mossy-nest swiftlet (*Aerodramus salangana*), Black-nest swiftlet (*Aerodramus maxima*), and Plume-toed swiftlet (*Collocalia esculenta*).

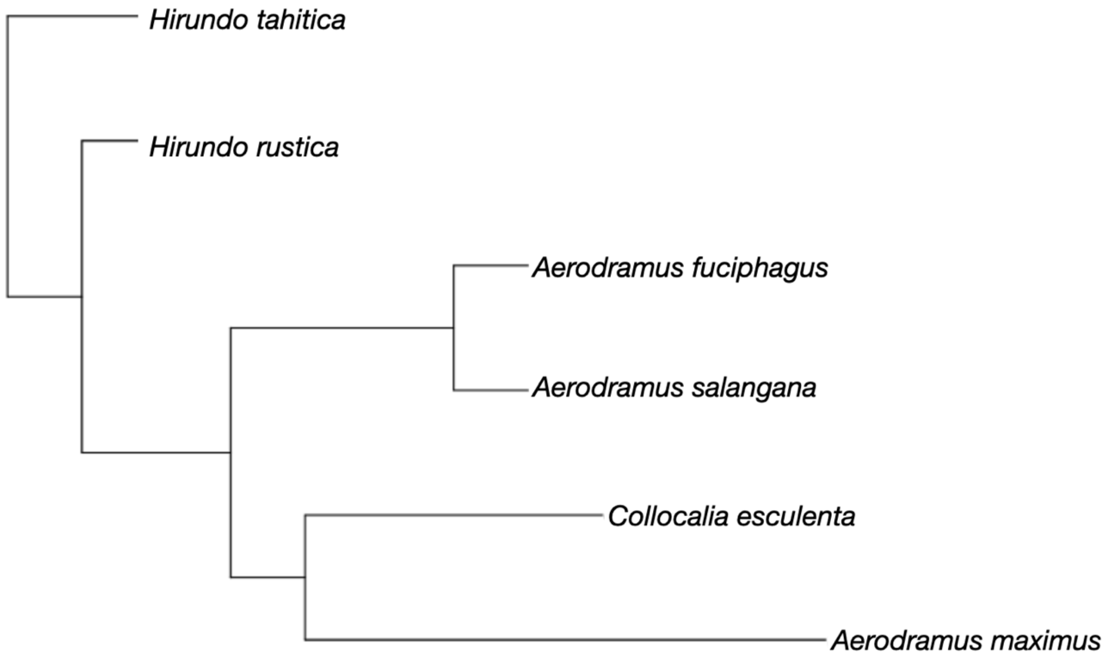

**Figure S2.** *Geographic niche overlap among environmental predictor sets.* Models incorporating human land-use variables alone predicted significantly greater geographic niche overlap among species compared to models using climate-only or combined predictors (Tukey post-hoc: Human land-use only - Climate-only, $p = 0.001$; Human land-use only - Both predictor sets, $p = 0.008$). No difference was detected between climate-only and combined models. Boxes show the interquartile range, horizontal lines show medians, and points represent pairwise species comparisons ($n = 15$ per predictor set).

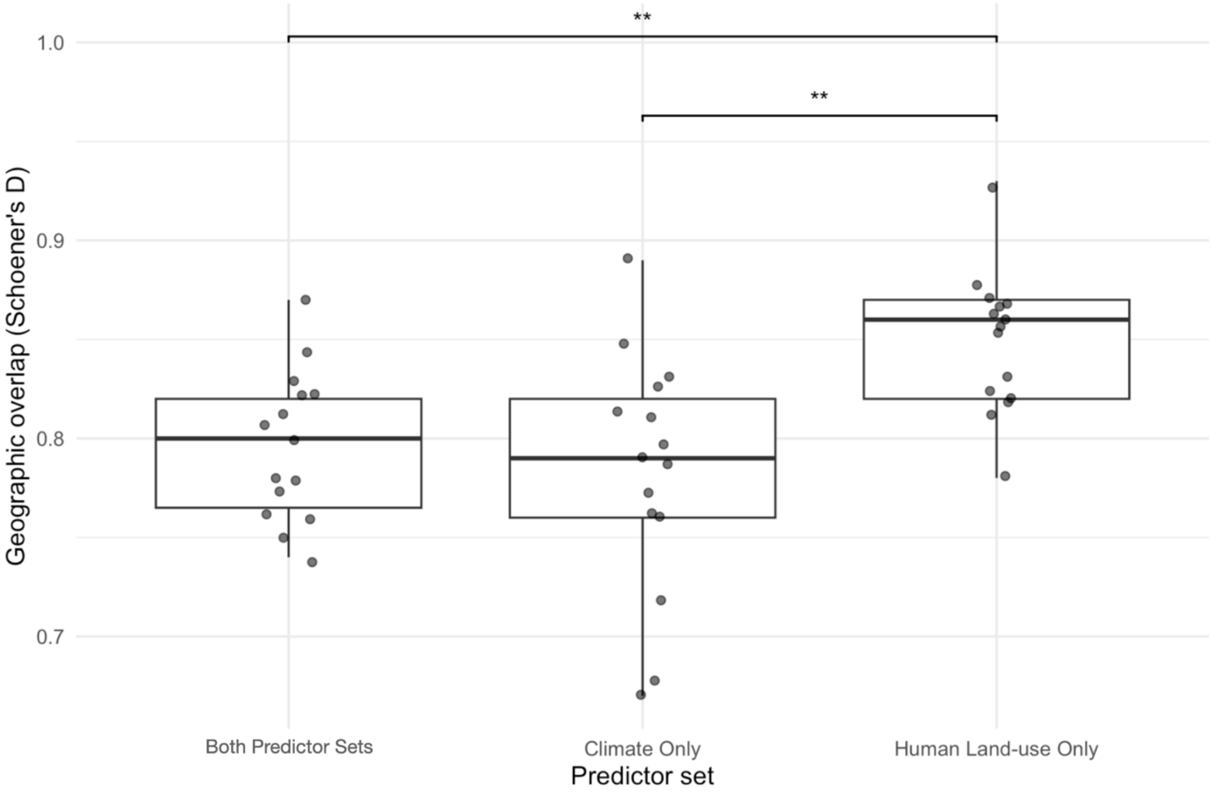

**Figure S3 –** Model 1 potential scale reduction factors (PSRF) for Beta, Omega, and Gamma parameters (morphology) indicate adequate MCMC convergence, with most values at or below 1.2.

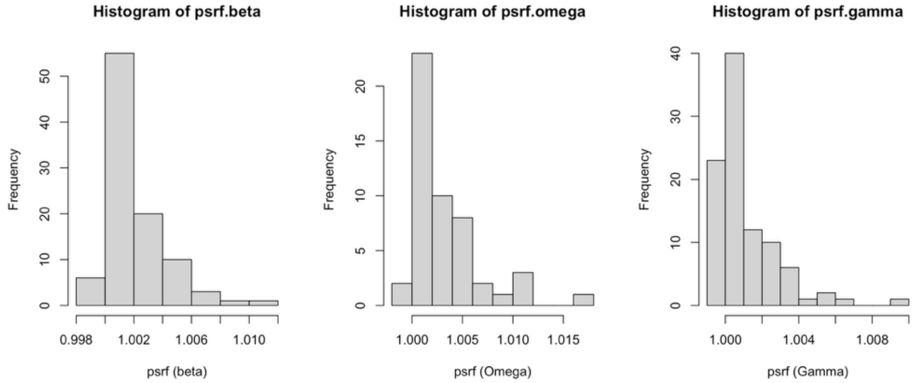

**Figure S4** – Model 2 potential scale reduction factors (PSRF) for Beta, Omega, and Gamma parameters (morphology and behavior) indicate adequate MCMC convergence, with most values at or below 1.2.

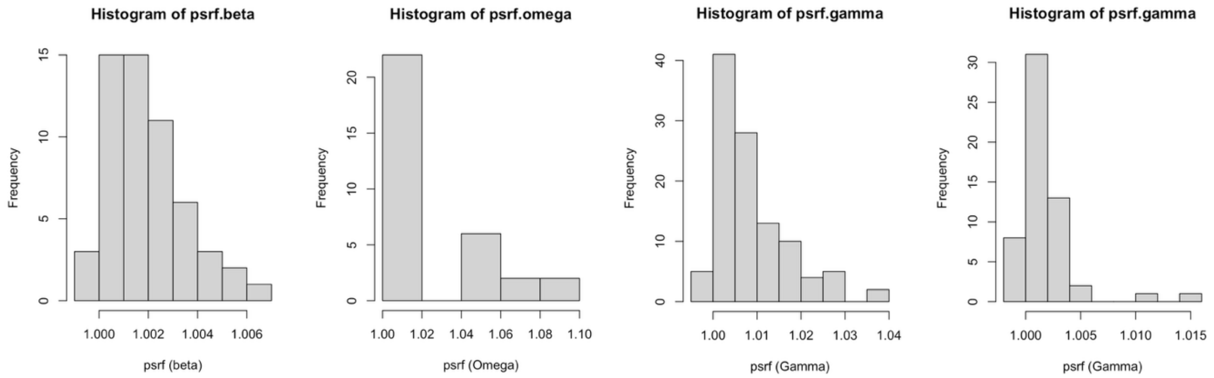

**Figure S5** – Model 3 potential scale reduction factors (PSRF) for Beta, Omega, and Gamma parameters (morphology and behavior) indicate adequate MCMC convergence, with most values at or below 1.2.

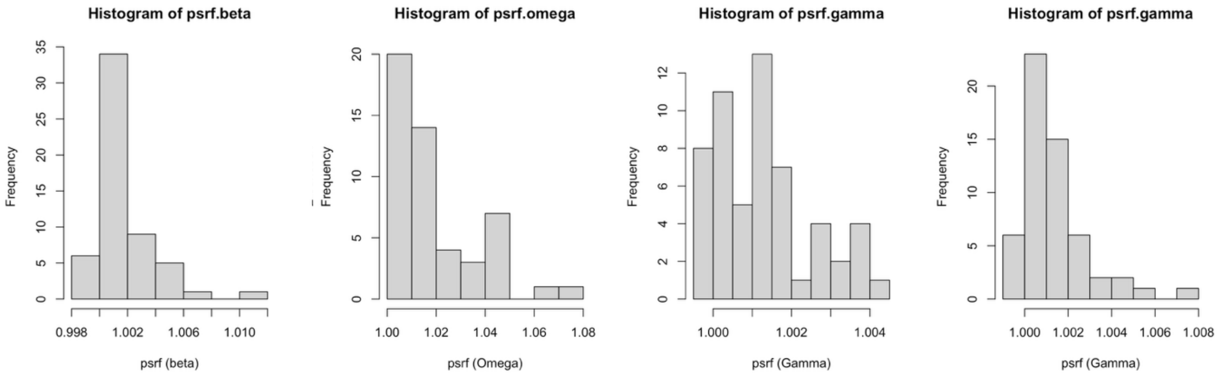

**Figure S6** – Model 4 potential scale reduction factors (PSRF) for Beta, Omega, and Gamma parameters (morphology and behavior) indicate adequate MCMC convergence, with most values at or below 1.2.

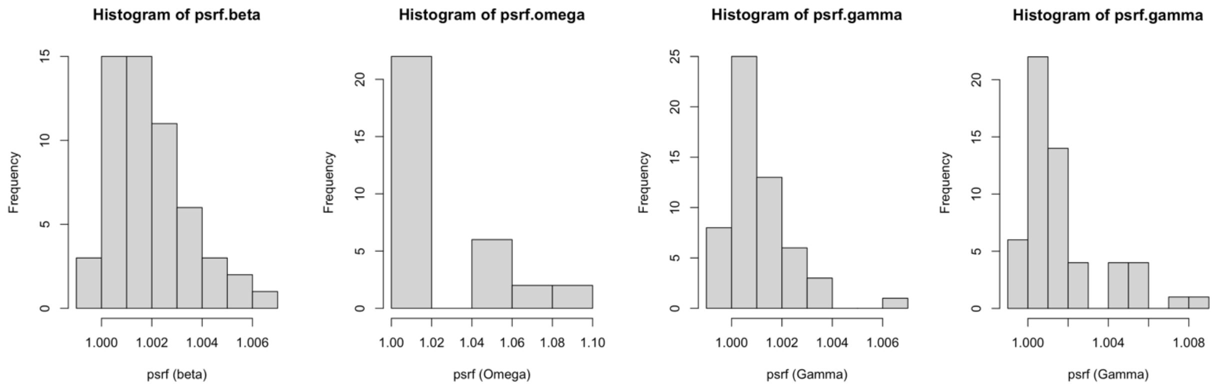

**Figure S7.** *Posterior Γ estimates for morphological trait–environment relationships across JSDMs.* Rows represent environmental predictors, and columns represent functional traits. Estimates reflect the posterior means of the gamma parameters, which describe how trait values influence species' responses to each environmental variable. Positive associations (in pink) indicate that species with higher trait values are more likely to occur in areas with higher values of that environmental variable; negative associations (in green) indicate the opposite. Only associations with moderate (posterior support 0.80–0.95) or strong (posterior support >0.95) support are shown.

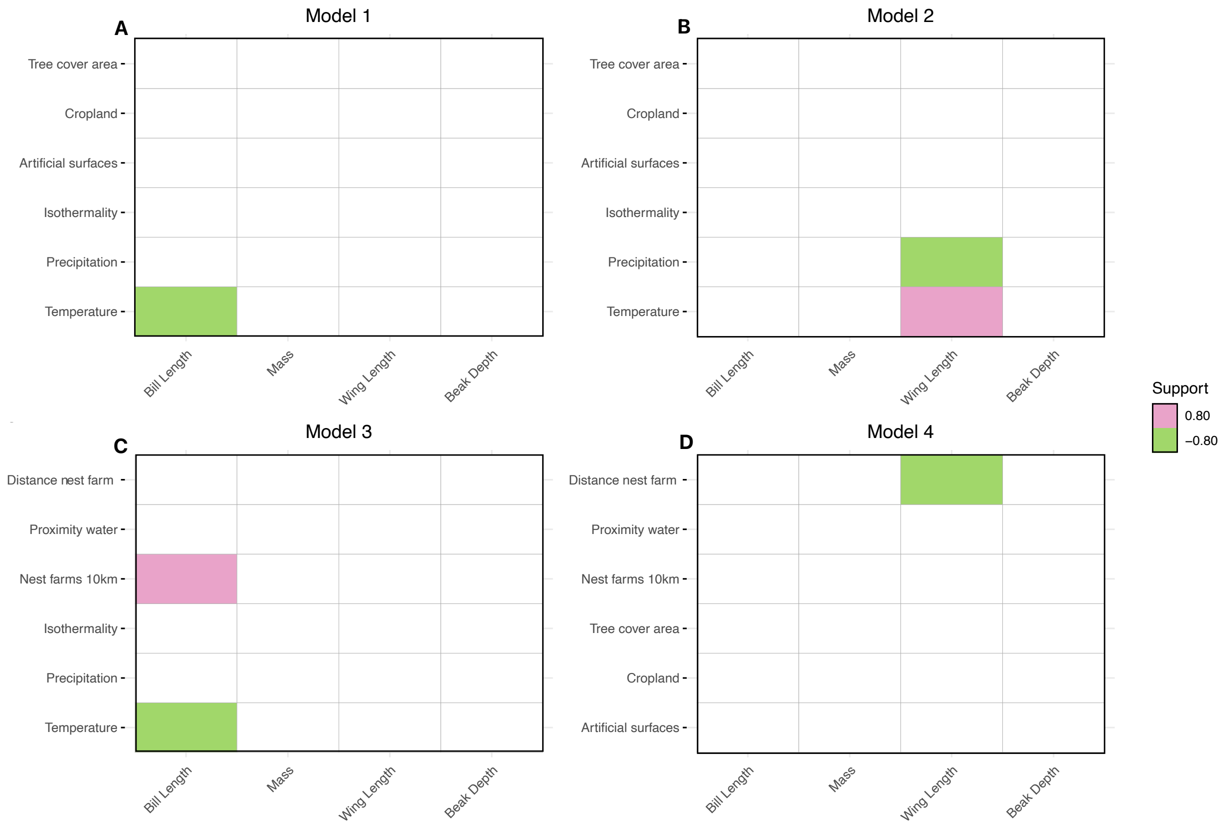

**Figure S8.** *Posterior Γ estimates for behavioral trait–environment relationships across three JSDMs (Models 2–4).* Rows represent environmental predictors, and columns represent functional traits. Estimates reflect the posterior means of the gamma parameters, which describe how trait values influence species' responses to each environmental variable. Positive associations (in pink) indicate that species with higher trait values are more likely to occur in areas with higher values of that environmental variable; negative associations (in green) indicate the opposite. Only associations with moderate (posterior support 0.80–0.95) or strong (posterior support >0.95) support are shown.

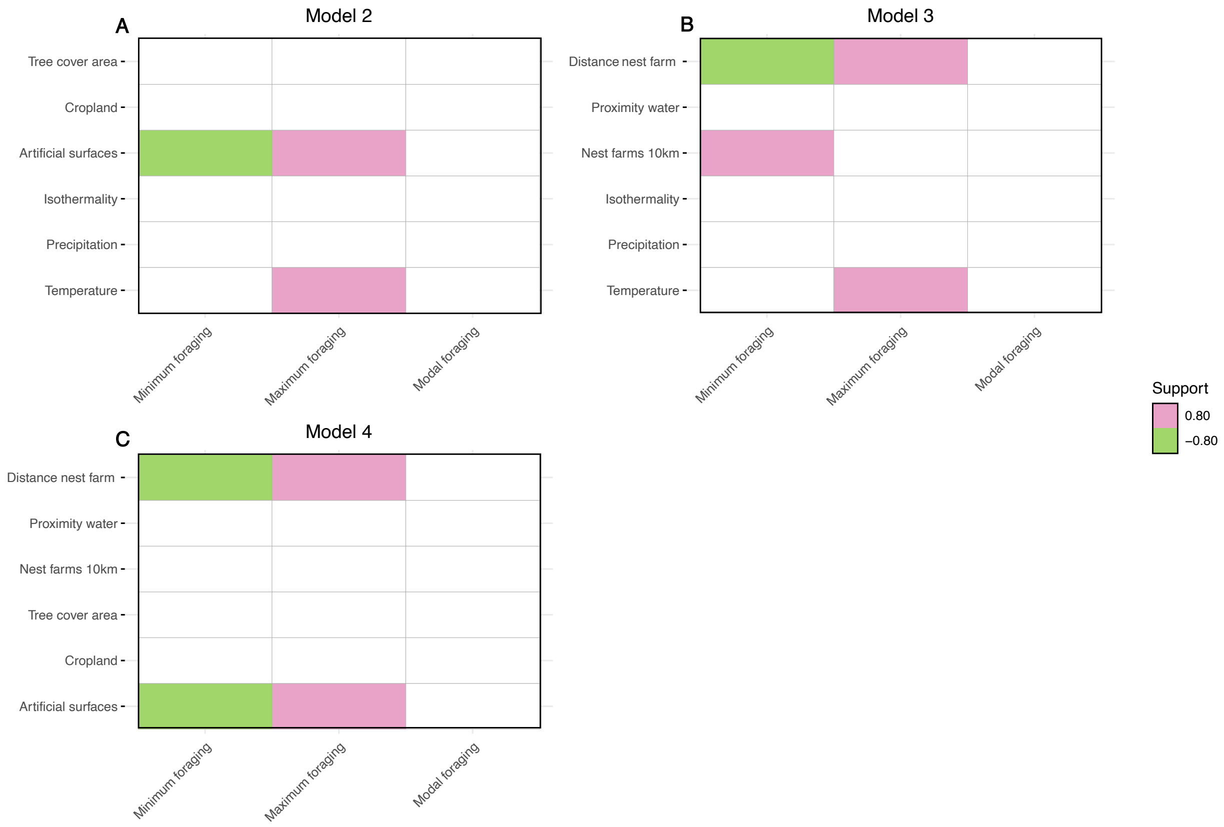
